# The PTEX pore component EXP2 is important for intrahepatic development during the *Plasmodium* liver stage

**DOI:** 10.1101/2022.10.02.510306

**Authors:** Tahir Hussain, Jose Linera-Gonzalez, John M Beck, Manuel A Fierro, Gunnar R Mair, Ryan C Smith, Josh R Beck

## Abstract

During vertebrate infection, obligate intracellular malaria parasites develop within a parasitophorous vacuole which constitutes the interface between the parasite and its hepatocyte or erythrocyte host cells. To transcend this barrier, *Plasmodium* spp. utilize a dual-function pore formed by EXP2 for nutrient transport and, in the context of the PTEX translocon, effector protein export across the vacuole membrane. While critical to blood stage survival, less is known about EXP2/PTEX function in the liver stage, although major differences in the export mechanism are indicated by absence of the PTEX unfoldase HSP101 in the intrahepatic vacuole. Here, we employed the glucosamine-activated *glmS* ribozyme to study the role of EXP2 during *Plasmodium berghei* liver stage development in hepatoma cells. Insertion of the *glmS* sequence into the *exp2* 3’UTR enabled glucosamine-dependent depletion of EXP2 after hepatocyte invasion, allowing separation of EXP2 function during intrahepatic development from a recently reported role in hepatocyte invasion. Post-invasion EXP2 knockdown reduced parasite size and largely abolished expression of the mid to late liver stage marker LISP2. As an orthogonal approach to monitor development, EXP2-*glmS* parasites and controls were engineered to express nanoluciferase. Activation of *glmS* after invasion substantially decreased luminescence in hepatoma monolayers and in culture supernatants at later time points corresponding with merosome detachment that marks the culmination of liver stage development. Collectively, our findings extend the utility of the *glmS* ribozyme to study protein function in the liver stage and reveal EXP2 is important for intrahepatic parasite development, indicating PTEX components also function at the hepatocyte-parasite interface.

## Introduction

Malaria remains one of the most devastating parasitic diseases in the world. In 2020, 241 million new infections occurred, resulting in 627 thousand deaths, mostly among children under the age of five in sub-Saharan Africa (1). Humans are infected by the bite of an *Anopheline* mosquito that deposits sporozoites in the skin which then travel to the liver and invade hepatocytes. A single round of development within this intrahepatic niche generates thousands of erythrocyte-infective forms which are released into the circulation to initiate the pathogenic blood stage (2). As a critical bottleneck to patent *Plasmodium* infection in the mammalian host, the liver stage has emerged as an important drug and vaccine target.

Following invasion of both erythrocytes and hepatocytes, the parasite develops within in a parasitophorous vacuole (PV) membrane (PVM) that forms the boundary of a compartment for expansive growth and the direct interface for host-parasite interactions (3, 4). In the blood stage, an arsenal of effector proteins is delivered across the PVM, remodeling the host cell to create a niche for intracellular development and avoid host defenses (5, 6). Protein export into the erythrocyte is facilitated by the *Plasmodium* Translocon of EXported proteins (PTEX) (7-9). The PVM-spanning channel of the translocon is formed by an oligomer of EXP2, the only widely conserved component of PTEX (10-12). EXP2 and its orthologs mediate small molecule exchange between the host cytosol and PV lumen of vacuole-dwelling apicomplexans (13). However, *Plasmodium spp*. have additionally functionalized this nutrient-permeable channel through a flange-like adaptor formed by PTEX150 which docks the AAA+ chaperone unfoldase HSP101, transforming EXP2 into a protein-conducting PVM translocon (10, 11, 14). In this way, proteins destined for export are unfolded by HSP101 in the PV and passed through PTEX150 and EXP2 across the PVM and into the erythrocyte cytoplasm (11).

While small molecule transport and protein export across the PVM are also anticipated to be critical for survival within the hepatocyte, the function of EXP2/PTEX is less clear during the liver stage. Intriguingly, while EXP2 and PTEX150 are also expressed during hepatocyte infection, HSP101 has not been detected until late liver stage development when it is loaded into the dense granules of forming merozoites to be deployed upon red blood cell (RBC) invasion (15-17). Furthermore, blood-stage export reporters are not translocated beyond the liver-stage PVM, even when HSP101 is ectopically expressed at this stage, indicating a distinct export mechanism (15, 18, 19). Inactivation of *exp2* expression in sporozoites using the FLP/FRT stage-specific conditional knockdown system in *P. berghei* demonstrated a critical role for EXP2 in transition from the mosquito host to the vertebrate blood-stage (15). However, subsequent work surprisingly reported this to be the result an invasion defect in these EXP2-deficient sporozoites which could be rescued by exogenous supplementation with recombinant EXP2 or bacterial pore forming toxins, implicating extracellular secretion of EXP2 in hepatocyte invasion through host membrane wounding (20).

While these findings imply a remarkable distinct function in hepatocyte entry, EXP2 is also expected to play an important role in PVM transport and possibly in effector protein export in the liver stage by analogy to its function in blood stage (10, 14) as well as the role of EXP2 orthologs GRA17/23 in PVM permeability in *Toxoplasma* (13). However, the functional importance of EXP2 during intrahepatic development remains unclear as existing genetic strategies provide limited control of *exp2* knockdown timing in the liver stage. Here, we applied the *glmS* ribozyme to study the role of EXP2 during liver stage parasite development, showing for the first time that this system can be used to modulate parasite gene expression beyond the blood stage. This strategy enables temporal control of knockdown, revealing that EXP2 plays a critical role in intrahepatic development of liver-stage parasites independent of any contribution to invasion, consistent with a function in PVM nutrient uptake and/or protein export into the hepatocyte.

## Results

### The *glmS* ribozyme enables inducible protein knockdown in the *P. berghei* blood and liver stages

To study EXP2 function in *Plasmodium* liver stage development after hepatocyte invasion, we sought to develop a ligand-based conditional knockdown approach that would enable more precise control of the timing of knockdown. We choose the *glmS* strategy which involves introduction of a bacterial metabolite-responsive ribozyme sequence immediately downstream of the stop codon of the target gene (21-23). Knockdown is initiated by addition of glucosamine (GlcN) to the culture medium, which upon conversion to glucosamine-6-phosphate, activates the ribozyme resulting in cleavage of the mRNA 3’ UTR and transcript destabilization, reducing target protein levels. The *glmS* ribozyme has been widely used to study protein function in the *P. falciparum* blood stage where the ability to indefinitely culture the parasite *in vitro* allows GlcN to be delivered at sufficient concentration and duration to mediate robust knockdown (23, 24). Indeed, the *glmS* approach is capable of producing lethal EXP2 knockdown in the *P. falciparum* blood stage (14). The *glmS* system has also been shown to function in the *P. berghei* blood stage where ∼2 mM GlcN was required to reduce the expression of a GFP reporter by 50% in culture, although difficulties in maintaining rodent malaria parasites *ex vivo* beyond a single developmental cycle limit the applicability of knockdown strategies requiring sustained presence of a ligand (25). Baseline GlcN serum concentrations are in the low nanomolar range in mice and can be increased to only ∼2 μM by dietary supplementation (26), still far below the range observed to activate substantial *glmS* cleavage in *Plasmodium* (23, 25). Thus while sufficient GlcN concentrations are unachievable for *in vivo* studies with *glmS*, endogenous GlcN levels are likely too low to significantly active the ribozyme in the rodent host so that tagging important or essential *P. berghei* genes with *glmS* should be tolerated.

To evaluate whether *glmS* can control *P. berghei* EXP2 levels, we first introduced an mRuby3-3xHA tag followed by the *glmS* sequence at the endogenous *P. berghei exp2* locus by double homologous recombination (Fig. 1A and S1). The donor plasmid also contains a cassette for expression of GFP driven by the constitutive *Pbhsp70* promoter. In parallel, we also generated a control line that was identical except for lacking the *glmS* sequence. Addition of GlcN to *ex vivo* cultures for 18 hours substantially reduced EXP2 levels relative to untreated controls in the *glmS* line but did not reduce EXP2 levels in the control parasites lacking the ribozyme (Fig. 1B,C). Specifically, treatment with 0.5 mM GlcN reduced EXP2 levels by 23±12% while higher concentrations achieved ∼50% knockdown (57±12% or 46±12% at 1 or 2 mM GlcN, respectively) (Fig. 1C), similar to previous reports (25). These results indicate the *glmS* ribozyme can modulate *P. berghei* EXP2 levels although blood stage knockdown is less substantial than that observed by *glmS* in *P. falciparum* (14), likely due to the inability to initiate GlcN treatment in the preceding cycle of *P. berghei* development. Furthermore, parasites used to initiate *ex vivo* cultures were not synchronized and thus any cells beyond ring-stage development will have already expressed substantial EXP2 prior to ribozyme activation (10).

**Figure 1.**
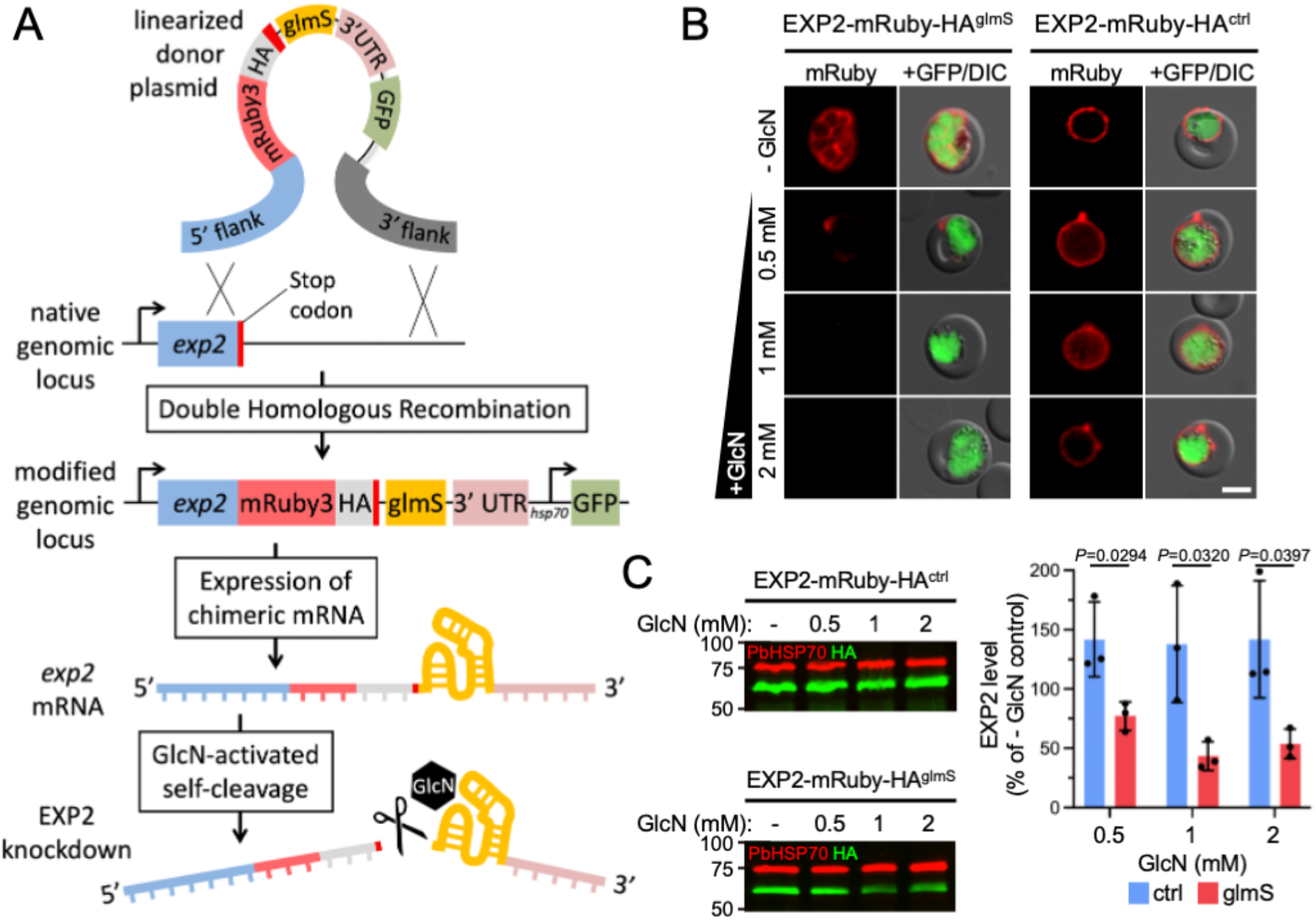
The *glmS* ribozyme mediates EXP2 knockdown in the *P. berghei* blood stage. A) Schematic showing strategy for double homologous recombination to install an mRuby3-3xHA tag followed by the *glmS* sequence immediately downstream of the stop codon at the endogenous *exp2* locus (EXP2-mRuby-HA^glmS^). The construct also contains a downstream cassette for expression of cytosolic GFP. Following transcription, the ribozyme is activated by binding to GlcN-6-phosphate, resulting in autocleavage to remove the mRNA 3’ UTR. A control line, identical except for lacking the *glmS* sequence, was also generated (EXP2-mRuby-HA^ctrl^, not shown). B) Live fluorescence imaging and C) Western blot of EXP2-mRuby-HA^glmS^ and EXP2-mRuby-HA^ctrl^ parasites following 18 hours of *ex vivo* culture in the absence or presence of the indicated GlcN concentrations. PbHSP70 serves as a loading control. Molecular weights are predicted to be 75 kDa for PbHSP70 and 58.9 kDa for EXP2-mRuby-HA after signal peptide cleavage. Scale bars = 5μm. Results are representative of three independent experiments. Quantification of EXP2-mRuby-HA levels are shown. Error bars indicate SD. P values were determined by an unpaired, two-sided Student’s t-test.

To determine the suitability of the *glmS* system for protein knockdown in cultured liver-stage parasites, we first evaluated whether GlcN levels sufficient to mediate knockdown in blood-stage parasites had any effect on proliferation of Huh7 or HepG2 hepatoma cell lines commonly used to cultivate the *P. berghei* liver stage (27). We assessed viability of hepatoma cells by quantifying metabolic activity using a resazurin assay (28) and found that proliferation of both Huh7 and HepG2 cell types was not reduced at GlcN concentrations up to 1 mM (Fig. S2). These results indicate GlcN can be supplied in hepatoma cultures at concentrations suitable for maximal blood-stage knockdown without host cell toxicity. As both HepG2 and Huh7 were similarly tolerant to GlcN levels, Huh7 cells were chosen for subsequent experiments due to their superior qualities for imaging intracellular parasites.

To evaluate whether the *glmS* ribozyme can mediate control of parasite protein levels in the liver stage, we next generated a line bearing a 3xFLAG epitope tag followed by the *glmS* sequence (or not as a control) at the endogenous *P. berghei exp2* locus. This plasmid also contains a downstream cassette expressing nanoluciferase (NanoLuc) under the control of the *Pbhsp70* promoter to provide a sensitive proxy for monitoring parasite development (29). Clonal lines of each strain were derived and designated EXP2^glmS^ and EXP2^ctrl^, respectively (Fig. S3). These lines showed similar levels of exflagellation and following blood feeding to mosquitoes, both lines produced similar numbers of sporozoites per mosquito as parental parasites, indicating that modification of the *exp2* genetic locus did not impact transmission to or development in the insect vector (Fig. S4).

EXP2^glmS^ and EXP2^ctrl^ sporozoites were isolated by mosquito dissection and allowed to infect Huh7 cells and develop in the presence or absence of 0.3 mM, 0.5 mM or 1 mM GlcN for 48 hours before cells were fixed and processed for immunofluorescence. In untreated EXP2^glmS^ cultures, EXP2 was readily observed at the parasite periphery where it largely co-localized with the liver-stage integral PVM marker UIS4 (30) (Fig. 2A, top panels). In contrast, EXP2^glmS^ parasites grown with GlcN displayed a striking reduction in EXP2 intensity by ?73-87% while UIS4 levels were not affected (Fig. 2A,B). Importantly, EXP2 levels in the control line were not similarly altered by GlcN, indicating the reduction observed in the EXP2^glmS^ parasites was a result of the ribozyme sequence in the *exp2* 3’ UTR and not an indirect effect of GlcN treatment (Fig. 2C,D). Notably, introduction of 0.5 mM GlcN at the time EXP2^glmS^ sporozoites were added to Huh7 host cells (0 hpi) had no impact on invasion (Fig S5) but reduced EXP2 levels by ∼80% at 48 hpi when maintained in cultures (Fig 2B). Moreover, similar EXP2 knockdown was achieved when GlcN was added after sporozoite invasion had occurred (3 hpi, Fig. 2A,B), clearly indicating successful EXP2 depletion after host cell entry without confounding effects from perturbed invasion. Collectively, these results show that the *glmS* ribozyme can be activated in the liver stage, enabling temporal control of parasite protein knockdown.

**Figure 2.**
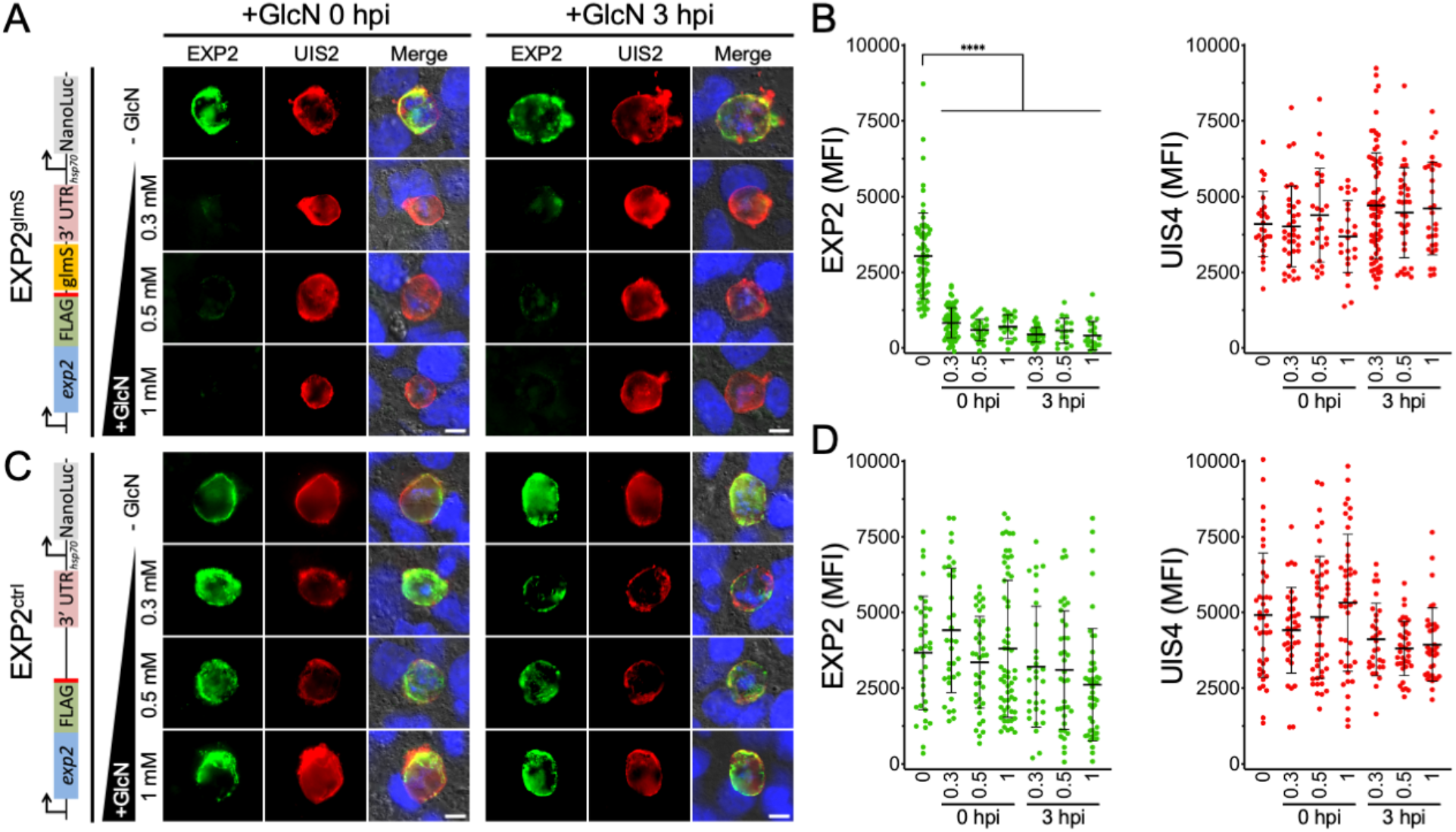
The *glmS* ribozyme mediates EXP2 knockdown in the *P. berghei* liver stage. A) Immunofluorescence assay on EXP2-FLAG^glmS^ liver-stage parasites after 48 hours of development in Huh7 cells detecting FLAG (EXP2) in green and UIS4 in red. GlcN was added at indicated concentrations at the time sporozoites were introduced (0 hpi, left panels) or 3 hours later after sporozoites had invaded (3 hpi, right panels). Merged panels also show DAPI in blue and DIC. Schematic at left shows modified *exp2* locus. Images are representative of three independent experiments. Scale bars = 5 μm. B) Quantification of EXP2 and UIS4 intensity. Data are pooled from three independent experiments. Error bars represent SD. C) Immunofluorescence assay on EXP2-FLAG^ctrl^ liver-stage parasites and D) quantification as in (B). Significance was determined by an unpaired, two-sided Student’s t-test. **** indicates *p*<0.0001.

### EXP2 knockdown reduces liver-stage parasite size and severely impacts intrahepatic parasite development

To determine the impact of EXP2 knockdown on intrahepatic parasite development, EXP2^ctrl^ and EXP2^glmS^ parasites were allowed to develop in Huh7 cells for 48 hours in the presence or absence of 0.3, 0.5 or 1 mM GlcN and vacuole size was determined by automated microscopy using UIS4 as a marker for the PVM (Fig. S6). While EXP2^ctrl^ parasite size was not altered by GlcN treatment (Fig. 3A), EXP2^glmS^ parasites showed a significant decrease in vacuole area at all GlcN treatment levels with mean vacuole size reduced by as much as 36% (Fig. 3B, +1 mM GlcN at 0 hpi). A similar decrease in parasite size was observed when GlcN was not added until after sporozoite invasion had occurred (Fig. 3B, +GlcN 3 hpi).

**Figure 3.**
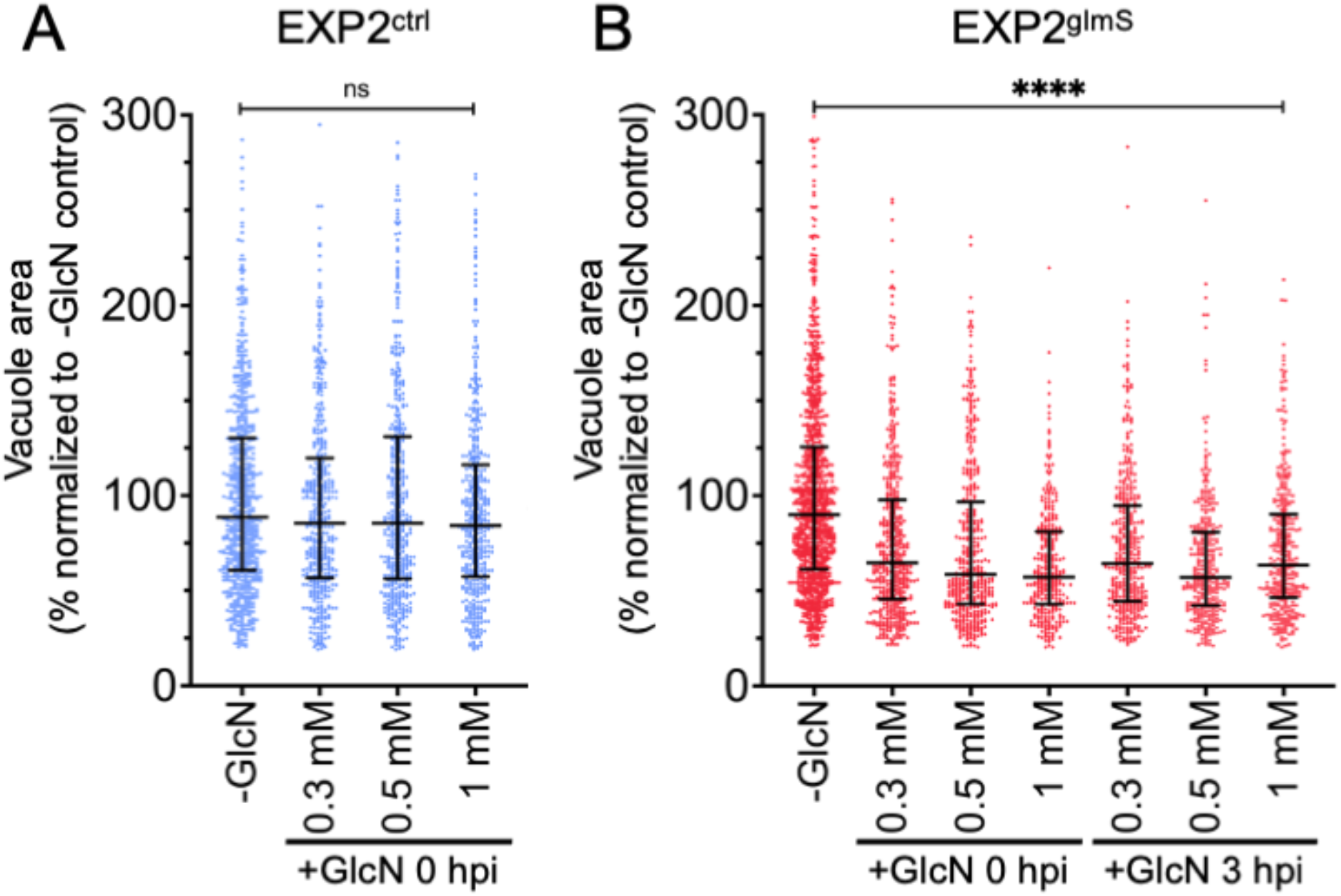
EXP2 knockdown reduces liver-stage parasite size. Quantification of parasite vacuolar area at 48 hpi in A) EXP2^ctrl^ or B) EXP2^glmS^. GlcN was introduced to cultures at the indicated concentrations and times (0 or 3 hpi). Data are pooled from three independent experiments and expressed as percentage of the mean vacuole area in the untreated controls. Median and interquartile ranges are shown. P values were determined by a Kruskal-Wallis test. ****, P<0.0001.

To provide an orthogonal, ultrasensitive measurement of parasite development, we next monitored luminescence generated from the constitutive NanoLuc expression cassette installed downstream of *exp2* (Fig. S3A). Parasites were again allowed to develop 48 hrs in the presence or absence of 0.3, 0.5 or 1 mM GlcN before measuring luminescence of the infected monolayer. As a control for complete parasite death, we also included an atovaquone treatment (31). Luminesce in EXP2^glmS^ cultures exposed to GlcN was strikingly reduced by as much as 78.4±9.4% (+1 mM GlcN 3 hpi) but did not substantially impact NanoLuc signal in EXP2^ctrl^ parasites (Fig. 4). Importantly, similar reduction in EXP2^glmS^ luminescence was observed regardless of whether GlcN was added at 0 or 3 hpi, indicating this effect was not the result of a reduction in sporozoite invasion.

**Figure 4.**
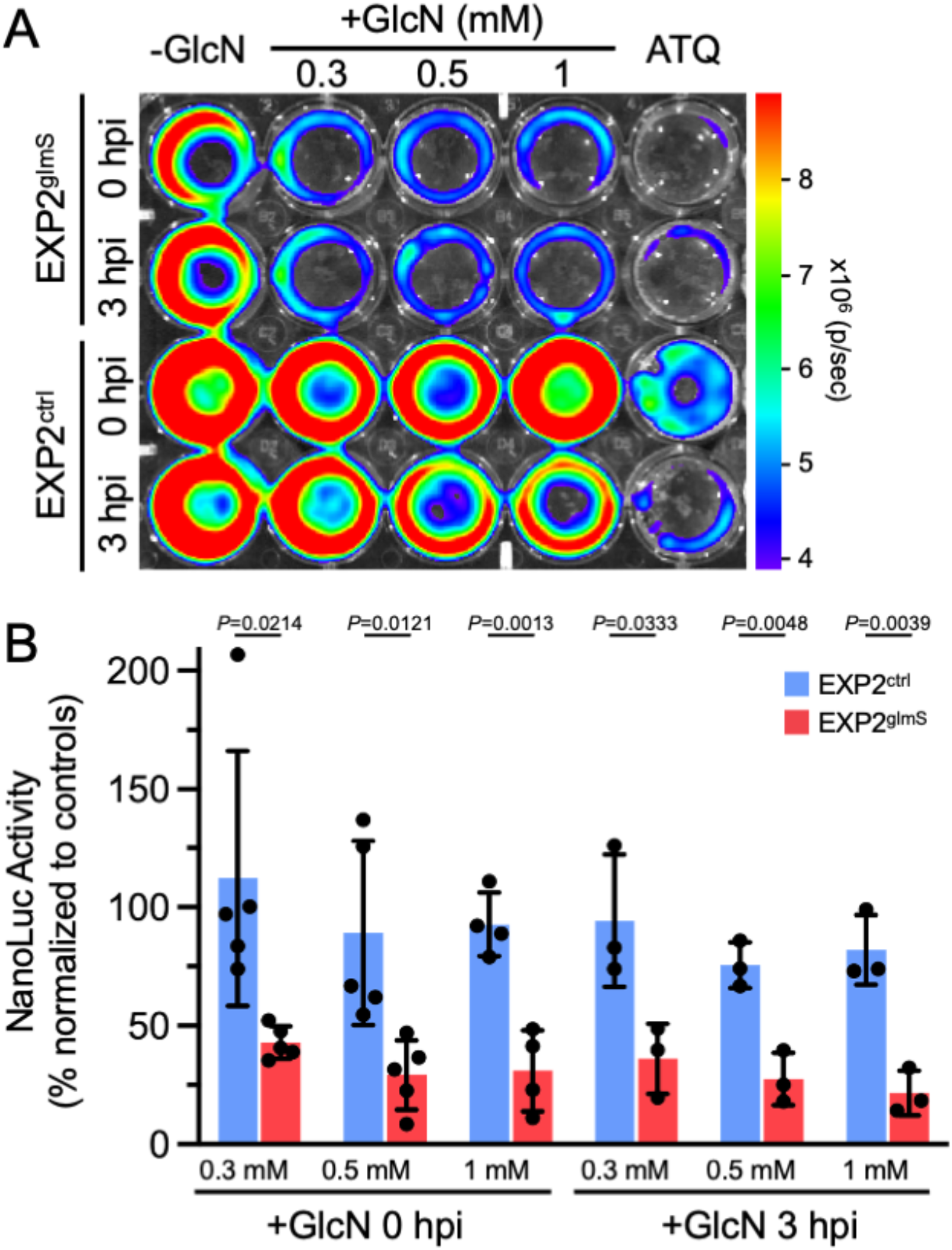
EXP2 knockdown severely impacts a liver-stage bioluminescence developmental reporter. A) Representative IVIS image of NanoLuc luminescence in Huh7 cultures 48 hpi with EXP2^glmS^ and EXP2^ctrl^. GlcN was introduced to cultures at the indicated concentrations and times (0 or 3 hpi). Atovaquone (ATQ, 10 nM) was added at 0 hpi as a control for parasite death. B) Quantification of luminescence in EXP2^glmS^ and EXP2^ctrl^ (red and blue bars, respectively). Data were normalized to untreated (-GlcN) and ATQ controls (used to define 100% and 0% NanoLuc activity, respectively, in each experiment). n=3-5 independent biological replicates. Error bars indicate SD. P values were determined by an unpaired, two-sided Student’s t-test.

Liver-stage development culminates in detachment of infected cells from the liver matrix as merosomes which enter the circulation to initiate the blood stage, a process that is recapitulated in culture by detachment of merosomes from the monolayer and release into the media (29, 32, 33). To determine if parasites depleted of EXP2 were able to complete the liver-stage and form merosomes, we measured NanoLuc signal in culture supernatants at 65 hpi. While luminescence was not impacted by GlcN treatment in EXP2^ctrl^ cultures, indicating productive merosome release, EXP2^glmS^ parasites showed a severe reduction in supernatant luminescence by as much as 74.9±12.5% (+1 mM GlcN), indicating that loss of EXP2 has a major impact on successful completion of the liver stage (Fig. 5). Collectively, these results show that EXP2 is important for intrahepatic parasite development independent of any contribution to host cell entry.

**Figure 5.**
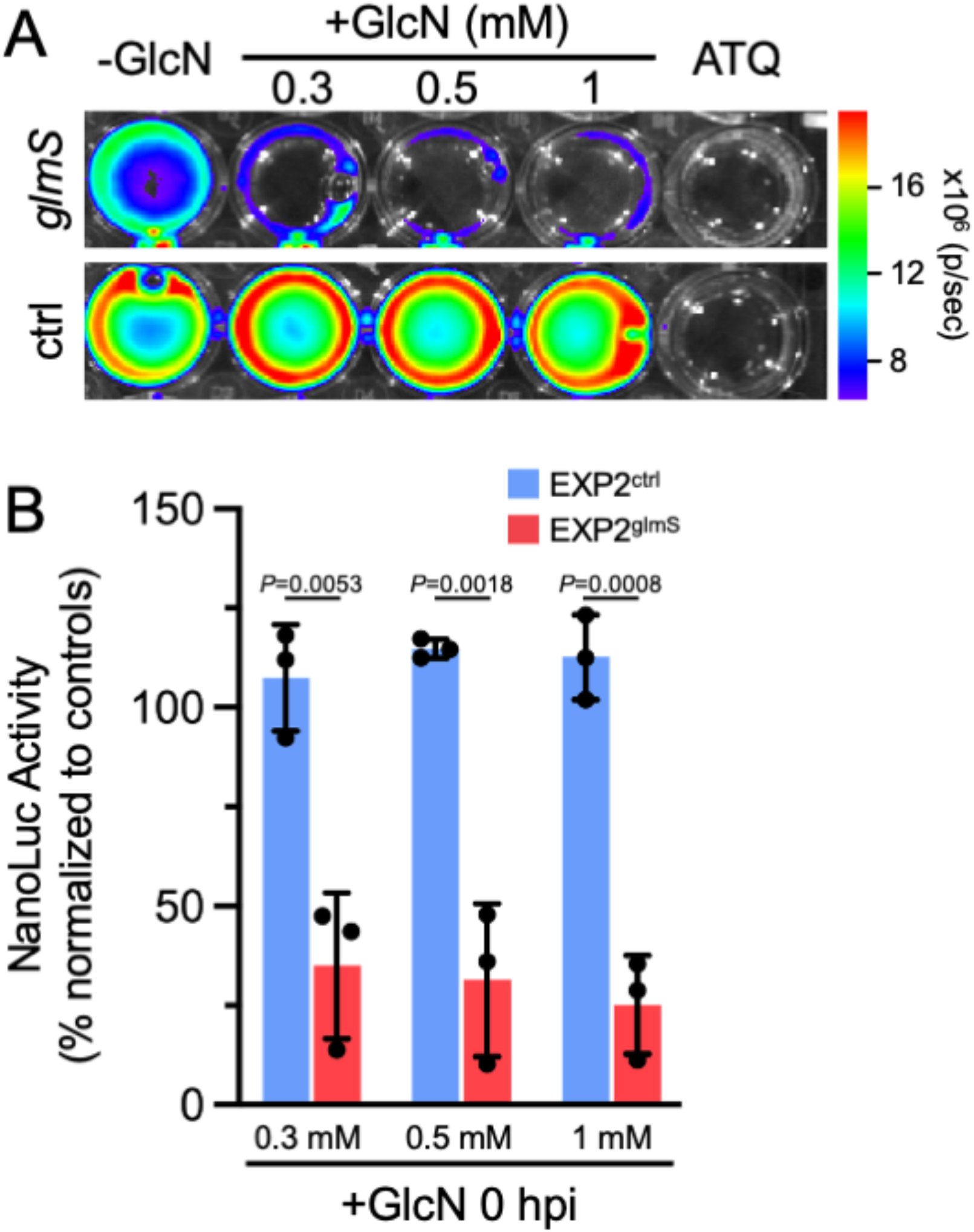
EXP2 knockdown reduces the formation of detached merosomes. A) Representative IVIS image of NanoLuc luminescence in Huh7 culture supernatants 65 hpi with EXP2^glmS^ and EXP2^ctrl^. GlcN was introduced to cultures at the indicated concentrations at 0 hpi. Atovaquone (ATQ, 10 nM) was added at 0 hpi as a control for parasite death. B) Quantification of luminescence in EXP2^glmS^ and EXP2^ctrl^ (red and blue bars, respectively). Data were normalized to untreated (-GlcN) and ATQ controls (used to define 100% and 0% NanoLuc activity, respectively, in each experiment). n=3 independent biological replicates. Error bars indicate SD. P values were determined by an unpaired, two-sided Student’s t-test.

### EXP2 knockdown decreases expression of the liver-stage developmental marker LISP2

To better understand when liver-stage defects occur following EXP2 knockdown, we examined expression of Liver-specific protein 2 (LISP2). LISP2 is expressed from the midpoint of development (beginning ∼24 hpi in *P. berghei*) through the end of schizogony, providing a marker for transition into the later stages of liver-stage development (34-36). We quantified LISP2 expression levels at 48 hpi in EXP2^glmS^ and EXP2^ctrl^ parasites with or without 0.3 or 0.5 mM GlcN treatment using LISP2 antisera raised against a peptide near the beginning of the C-terminal 6-Cys domain that localizes to the PV (34, 35). While LISP2 levels were not significantly altered by GlcN treatment in EXP2^ctrl^ parasites, EXP2^glmS^ parasites displayed a striking loss of LISP2 expression in the presence of GlcN with the majority of cells showing no apparent LISP2 signal (Fig. 6). Loss of LISP2 expression suggests that EXP2 is important for early liver-stage development and that parasites depleted of EXP2 do not proceed normally into schizogony, consistent with an anticipated function in nutrient uptake and/or protein export across the PVM.

**Figure 6.**
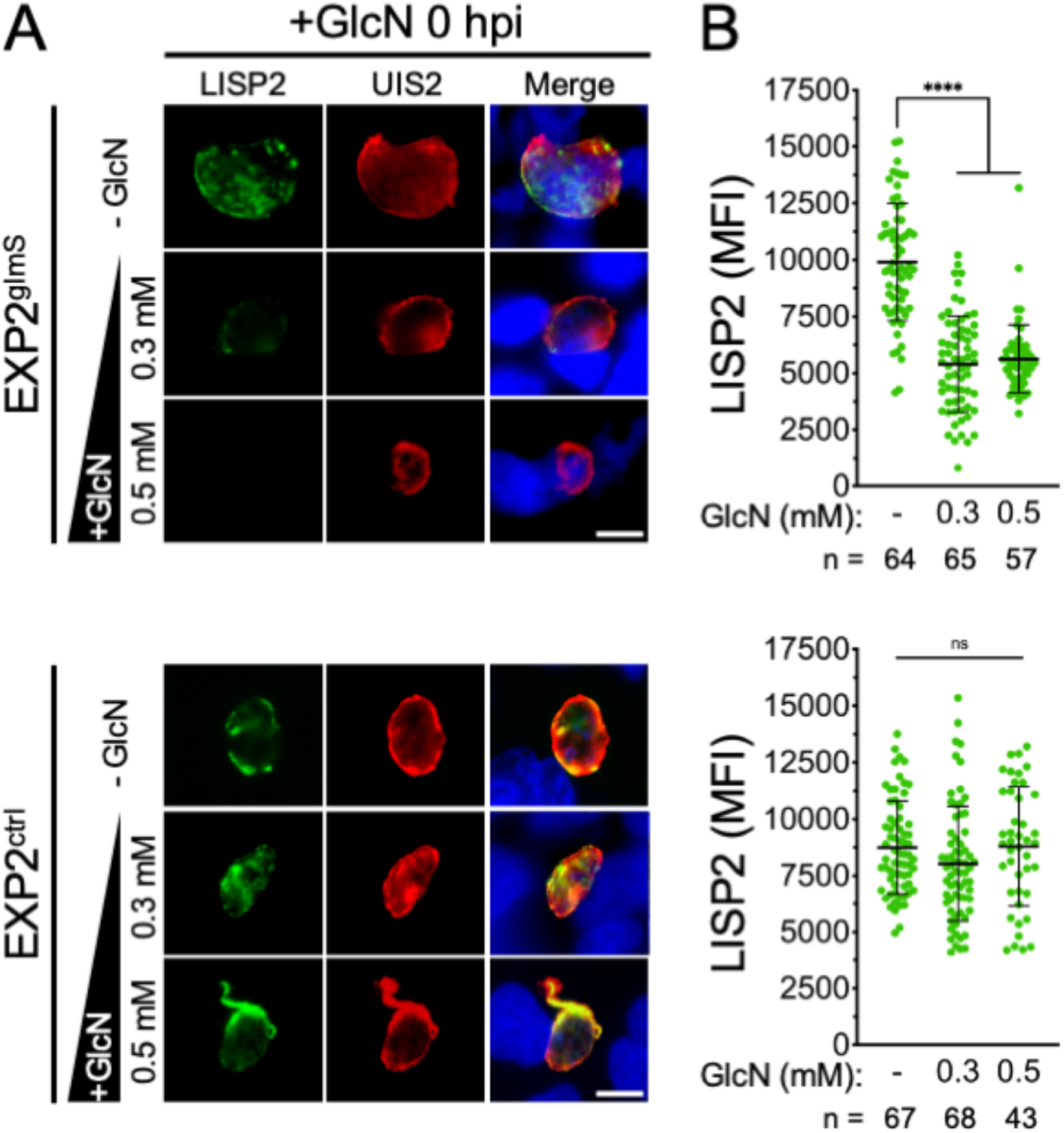
EXP2 knockdown abrogates expression of the mid to late stage liver-stage developmental marker LISP2. A) Immunofluorescence assay on EXP2-FLAG^glmS^ and EXP2-FLAG^ctrl^ parasites after 48 hours of development in Huh7 cells detecting LISP2 and UIS4. GlcN was added at indicated concentrations at the time sporozoites were introduced (0 hpi). Merged panels also show DAPI in blue. Scale bars = 5 μm. B) Quantification of LISP2 intensity. UIS4 was used to mark the PVM and LISP2 fluorescence within this area was collected. Data are pooled from three independent experiments. Error bars represent SD. Significance was determined by an unpaired, two-sided Student’s t-test. **** indicates P<0.0001.

## Discussion

Nutrient transport and effector protein translocation across the PVM are critical for the malaria parasite to establish its intracellular niche in the erythrocyte, avoid host defenses and fuel rapid parasite growth (37). Though less well understood, similar transport mechanisms are expected to be in operation at the hepatocyte-parasite interface during liver-stage infection. Indeed, the liver-stage vacuole is also permeable to small molecules (38) consistent with the presence of a similar nutrient-permeable channel as that observed in the blood stage and related apicomplexans (39-41). To promote their growth, liver-stage *Plasmodium* also recruit hepatocyte factors to the host-parasite interface, including the water/glycerol channel aquaporin 3 (42-44) and the endosomal-adaptor APPL1 (45). More broadly, parasites manipulate host gene expression (46, 47), inhibit apoptotic cell death (33) and remodel host cell biology (48) while avoiding hepatocyte cellular defenses such as vacuole-targeted selective autophagy (49, 50). Evasion of autophagy is mediated through sequestration of host LC3 by the PVM protein UIS3 (51) and other effectors secreted beyond the PV are likely to be involved in hepatocyte subversion.

While nutrient transport and protein export across the intraerythrocytic PVM both depend on the PTEX membrane pore EXP2 (10, 14), the intrahepatic function of EXP2/PTEX has remained unclear. A recent study unexpectedly reported that EXP2 is secreted by extracellular sporozoites and is important for hepatocyte invasion to initiate the liver stage (20). As EXP2 knockdown in the blood stage was not observed to impact erythrocyte invasion (10, 14), this additional EXP2 function is apparently unique to hepatocyte entry. However, whether EXP2 performs important roles in the PV during intrahepatic development has remained obscure. Here we show that the *glmS* ribozyme can be used to control parasite protein expression in the liver stage, enabling exploration of EXP2 function during intracellular development in hepatoma cells.

The ribozyme self-cleavage activity of *glmS* is predominantly activated by GlcN-6-P binding, and to lesser extent by unphosphorylated GlcN and related amine containing molecules (52). GlcN-6-P is generated from fructose-6-phosphase at the glycolytic branch point that feeds into the hexosamine biosynthesis pathway or by uptake and phosphorylation of GlcN from the environment. Differences in hexosamine metabolic flux between hepatoma and red blood cells notwithstanding, we found that basal Huh7 GlcN-6-P levels were not sufficient to produce EXP2 knockdown without addition of exogenous GlcN (Fig. 2B,D). In contrast, GlcN supplementation produced a robust and specific knockdown in EXP2^glmS^ parasites, reducing EXP2 levels by ∼75% and demonstrating ribozyme-mediated control of gene expression.

While similar GlcN concentrations activated EXP2 knockdown in blood- and liver-stage parasites, knockdown experiments in the blood-stage showed a more substantial dose-response to GlcN concentrations than the liver-stage knockdown. GlcN transport into mammalian cells occurs primarily through GLUT2, the major hexose transporter expressed in hepatocytes which has uniquely high affinity for GlcN over other substrates but also via GLUT4 or GLUT1, the predominant hexose transporter expressed in erythrocytes (53-56). Additionally, hepatoma cells show alterations in transporter expression and metabolism comparted to primary cells (57, 58) and GLUT1 activity is also increased in cells infected with liver-stage *P. berghei* (59). Thus, knockdown sensitivity could reflect differences in host cell GlcN uptake or transport across the parasite plasma membrane; alternatively, this might result from the longer incubation times and highly synchronous nature of the liver-stage experiments. To our knowledge, this is the first example of *glmS* utilization to control protein expression in an intracellular pathogen within a nucleated host cell, extending its range of application and providing a new tool to allow for temporal control of protein knockdown in the liver stage.

Invasion of EXP2^glmS^ sporozoites was not impacted when GlcN and parasites were introduced simultaneously into Huh7 cultures (Fig. S5), possibly due to insufficient time to achieve substantial mRNA degradation. In contrast, intrahepatic EXP2 levels could be depleted even when GlcN was not added until after sporozoite invasion. Analysis of vacuole size and NanoLuc reporter luminescence (as a proxy for parasite growth and maturation) clearly indicate EXP2 is important for intrahepatic development (Fig. 3-5). Furthermore, loss of LISP2 expression following EXP2 knockdown indicates a defect at an early stage, consistent with the established blood-stage roles of EXP2 in protein export and nutrient uptake that are critical to early parasite development in the erythrocyte (10, 14). In *Toxoplasma*, the EXP2 orthologs GRA17/23 function in PVM permeation but not in an analogous but mechanistically distinct PV protein export pathway; thus it is possible that EXP2 might function exclusively in small molecule transport during the liver stage (13, 37). The apparent absence of HSP101 in the liver-stage PV and the observation that several parasite proteins or reporters which are exported into the erythrocyte are not similarly translocated into the hepatocyte indicates that protein export is at least mechanistically distinct in the liver stage (15, 16, 18, 19). However, the upregulation PTEX150 in activated sporozoites (60) and its presence together with EXP2 in the liver-stage PV (17) suggests EXP2 is also functionalized for protein translocon in the hepatocyte, although it is unclear at present how this minimal translocon would be powered.

While hundreds of putative exported proteins have been identified in the blood stage, very little is known about the effector arsenal secreted into the hepatocyte with LISP2 being one of only a few parasite proteins reported to cross the PVM (34, 36). LISP2 is a large, conserved *Plasmodium* protein that contains a signal peptide and *bona fide* PEXEL motif followed by a repeat region and C-terminal 6-Cys domain (34, 61, 62). While the C-terminal portion of LISP2 detected by the antibody used in this study has not been observed to traffic beyond the PV, an N-terminal portion of the protein has been reported to be exported into the hepatocyte cytosol and nucleus, particularly in later stages of development (36 hpi or beyond in *P. berghei*) (34, 36). Unfortunately, the loss of LISP2 expression in parasites depleted of EXP2 prevented us from evaluating whether EXP2 was important for LISP2 translocation into the host cell. The identification of novel liver-stage exported proteins expressed early during infection along with determining the importance of PTEX150 to liver-stage development will help clarify the mechanism of protein export into the hepatocyte. Collectively, our findings extend the utility of the *glmS* ribozyme to study protein function in the liver stage and reveal an important function for EXP2 in intrahepatic parasite development, opening the door to studying liver-stage PVM transport mechanisms and their role in hepatocyte subversion.

## Materials and Methods

### Parasite maintenance and sporozoite production

*P. berghei* ANKA clone 2.34 and derivatives were maintained in Swiss Webster mice (Charles River). To enhance the production of parasite numbers, mosquito infections were performed in a TEP1 mutant strain (Δct1) of *Anopheles gambiae* (63, 64). Mosquitoes were reared at 27°C and 80% humidity on a 14/10 hour day/night cycle. Larvae were fed Tetramin (Tetra) and adults were maintained on 10% sucrose. For parasite infection, mosquitoes (4-6 days post-emergence) were fed on infected mice when parasitemia reach ∼5% parasitemia and exflagellation was observed. Blood fed mosquitoes were maintained on 10% sucrose at 21ºC for 21-28 days before isolation of salivary gland sporozoites. All experiments involving rodents were reviewed and approved by the Iowa State University Institutional Animal Care and Use Committee.

### Genetic modification of *P. berghei*

For generation of EXP2-mRuby3-3xHA *glmS* and control lines, 5’ and 3’ flanks targeting the 3’ end of *exp2* were amplified from parasite gDNA using primer pairs P1/2 and P3/4 and inserted between AvrII/XhoI and SacII/HpaI, respectively, in the plasmid pBAT1. The mRuby3-3xHA coding sequence was then amplified with P5/6 and inserted between HpaI/SphI. A 315 bp sequence containing the *glmS* ribozyme was amplified from plasmid pL6-3HA-glmS-BSD (65) with primers P7/8 and inserted at SphI between the stop codon and the downstream PbPPPK-DHPS 3’ UTR, resulting in the plasmid pBAT-EXP2-mRuby3-3xHA-*glmS*. To generate a matched control lacking only the ribozyme, the 186 bp comprising the *glmS* sequence were then removed using a QuickChange Lightning Multi Site Directed Mutagenesis kit (Aglient) and primer P9, resulting in the plasmid pBAT-EXP2-mRuby3-3xHA-control.

For generation of EXP2-3xFLAG *glmS* and control lines, sequence encoding a 3xFLAG tag was inserted between HpaI/SphI in pBAT-EXP2-mRuby3-3xHA-*glmS* using P10, replacing the mRuby3-3xHA. The NanoLuc coding sequence was then inserted between SwaI/BamHI using P11/12, replacing GFP and placing it under the control of the constitutive *Pbhsp70* promoter, resulting in the plasmid pBAT-EXP2-3xFLAG-*glmS*. To generate a matched control, the *glmS* sequence was then removed from this plasmid using a QuickChange Lightning Multi Site Directed Mutagenesis kit and primer P9 as above, resulting in the plasmid pBAT-EXP2-3xFLAG-control.

Plasmids were linearized at SacII/XhoI and transfected into *P. berghei* as described (66) except that schizont-infected RBCs were purified on an LD column mounted on a QuadroMACs magnetic separator (Miltenyi Biotech). Transfected parasites were reinjection into naïve mice and selection was applied with 0.07 mg/ml pyrimethamine in drinking water provided *ad libitum* 24 hours later. After returning from selection, transfected populations were cloned by limiting dilution IV injection into naïve mice to generate isogenic populations.

### GlcN treatment and evaluation of EXP2 knockdown

For blood-stage knockdown experiments, parasite-infected blood was collected by cardiac puncture and cultured in complete RPMI with or without GlcN supplementation. After 18 hours of culture, parasites were imaged by live fluorescence microscopy and collected for Western blot. For analysis of EXP2 knockdown by Western blot, parasites were harvested in cold PBS containing 0.035% saponin (Sigma), washed in PBS and lysed in RIPA buffer containing Halt protease inhibitor cocktail (Thermofisher). Lysates were briefly bath sonicated before centrifugation to pellet hemozoin. Cleared supernatants were lysed in sample buffer containing dithiothreitol and boiled before separation by SDS-PAGE. Primary antibodies were detected with IRDye 680- or 800-conjugated secondary antibodies (Li-COR Biosciences) used at 1:10,000. Western blot imaging was performed with an Odyssey infrared imaging system (Li-COR Biosciences) and signal quantification was carried out with Image Studio software (Li-COR Biosciences).

For liver-stage knockdown experiments, Huh7 cells were grown in DMEM media supplemented with 10% FBS and penicillin/streptomycin/glutamine (Gibco). Prior to infection, media was further supplemented with 2.5 μg/ml amphotericin B. Cells were seeded in 24-well plates or on coverslips and allowed to reach 60-80% confluency. To initiate infections, 150,000 freshly dissected sporozoites were centrifuged at 300xg for 5 min onto host cells and allowed to invade for 3 hours, after which media was replaced to remove uninfected sporozoites and subsequently change 3 times per day. GlcN was introduced at the time of sporozoite addition (0 hpi) or at the 3 hour media change (3 hpi).

For immunofluorescence (IFA) assays, cells were fixed with 4% paraformaldehyde for 15 minutes, permeabilized with PBS containing 0.1% Triton X-100 for 10 minutes and blocked in PBS containing 5% BSA for 1 hour. Blocked samples were incubated with primary antibodies diluted as indicated below for 1 hour followed by extensive washing. Primary antibodies were detected by incubation with Alexa Fluor 488 or 594 secondary IgG antibodies (Thermofisher) used at 1:2000 for 1 hour and following extensive washing, coverslips were mounted in ProLong Gold containing DAPI (Thermofisher). Live fluorescence and immunofluorescence images for display were collected with a 63x objective on an Axio Observer 7 equipped with an Axiocam 702 mono camera and Zen 2.6 Pro software (Zeiss) using the same exposure times for all images across sample groups and experimental replicates. For quantification of EXP2 expression and vacuolar area, images were collected using a 40x objective on a BZ-X810 automated microscope (Keyence). The thresholding tool was applied to the UIS4 channel to define a ROI corresponding to the boundary of each vacuole while investigators were blind to all other channels. The EXP2 and UIS4 or LISP2 mean fluorescence intensity within each ROI was then collected along with the area of each ROI. Vacuolar areas were normalized and plotted as percent of the mean area of untreated controls.

### Sporozoite invasion assays

EXP2^glmS^ sporozoites were allowed to infect Huh7 cells in the presence or absence of 0.5 mM GlcN as above. After 3 hours, media was changed before coverslips were fixed and processed for IFA. Images were acquired as above on a Keyence BZ-X810 automated microscope and invaded parasites were detected by thresholding UIS4 signal and area to distinguish newly formed vacuoles from uninvaded sporozoites to determine invaded parasites per sample area.

### Bioluminescence assays

Quantification of NanoLuc signal was carried out using a Nano-Glo Luciferase Assay Kit (Promega) according to the manufacturer’s recommendations. To monitor development in attached cells, 48 hpi liver-stage cultures were washed with PBS and lysed in Nano-Glo Luciferase Assay buffer containing Nano-Glo Luciferase Assay substrate diluted 1:200 and luminescence was measured after 3 minutes on an IVIS Specturum CT (PerkinElmer). To monitor merosome detachment at 65 hpi, parasite culture was carried out as above until 40 hpi when media was changed for a final time and the volume was reduced to 200 μl. At 65 hpi, media was collected and transferred to new plate and luminescence was recorded as above except that the Nano-Glo Luciferase Assay substrate was diluted 1:100 into the assay buffer. In each case, uninfected Huh7 cells were used as a negative control. To control for variance between independent sporozoite batches across biological replicates, luminescence signals were normalized to the untreated, 0 mM GlcN samples in each experiment. Data were then pooled and plotted as percent NanoLuc activity normalized to the mean luminescence in untreated, 0 mM GlcN controls (100%) and atovaquone treated controls (0%).

### Antibodies

The following antibodies were used for IFA and Western blot at the indicated dilutions: rabbit polyclonal anti-HA SG77 (Thermofisher; 1:1000 WB); mouse anti-PbHSP70 monoclonal antibody 4C9 (67) (WB 1:1000); mouse anti-FLAG monoclonal antibody M2 (Sigma; 1:1000 IFA); goat polyclonal anti-UIS4 (LS Biolabs LS-C204260; IFA 1:1000); rabbit polyclonal anti-LISP2 (IFA: 1:1000). Anti-LISP2 antisera was generated by Genscript against the peptide NGQKGNVDEERKSM located between the repeat region and 6-Cys domain, a portion of LISP2 previously shown to localize to the PV and but not observed to enter the host cell (35). Purified antisera labeled the parasite periphery in immunofluorescence assays and loss of signal following EXP2 knockdown provided validation of specificity.

### Resazurin assays

Huh7 or HepG2 cells were seeded at 4,000 cells/well in black-walled, clear-bottom 96 well plates and cultured in complete DMEM supplemented with a series of GlcN concentrations. After 72 hrs, 44 μM resazurin was added for 5 hours before recording fluorescence (Ex/Em 530/585) on a CLARIOstar Plus plate reader. Data were expressed as percent growth following normalization to minimum and maximum values. Dose-response curves were fit to the mean of four independent biological replicates and EC50 values were derived using Prism v9 (Graphpad).

## Acknowledgements

We thank F. Zavala for the PbHSP70 antibody. This study was supported by the Roy J. Carver Charitable Trust (http://www.carvertrust.org) grant 18-5102 to JRB. The funders had no role in study design, data collection and interpretation, or the decision to submit the work for publication.

## Author contributions

TH and JRB conceived and designed the experiments. TH, JL, JMB, GRM and JRB performed the experiments. RCS provided reagents. TH, MAF and JRB analyzed the data. TH and JRB wrote the manuscript. All authors discussed and edited the manuscript.

## Figure legends

**Figure S1.**
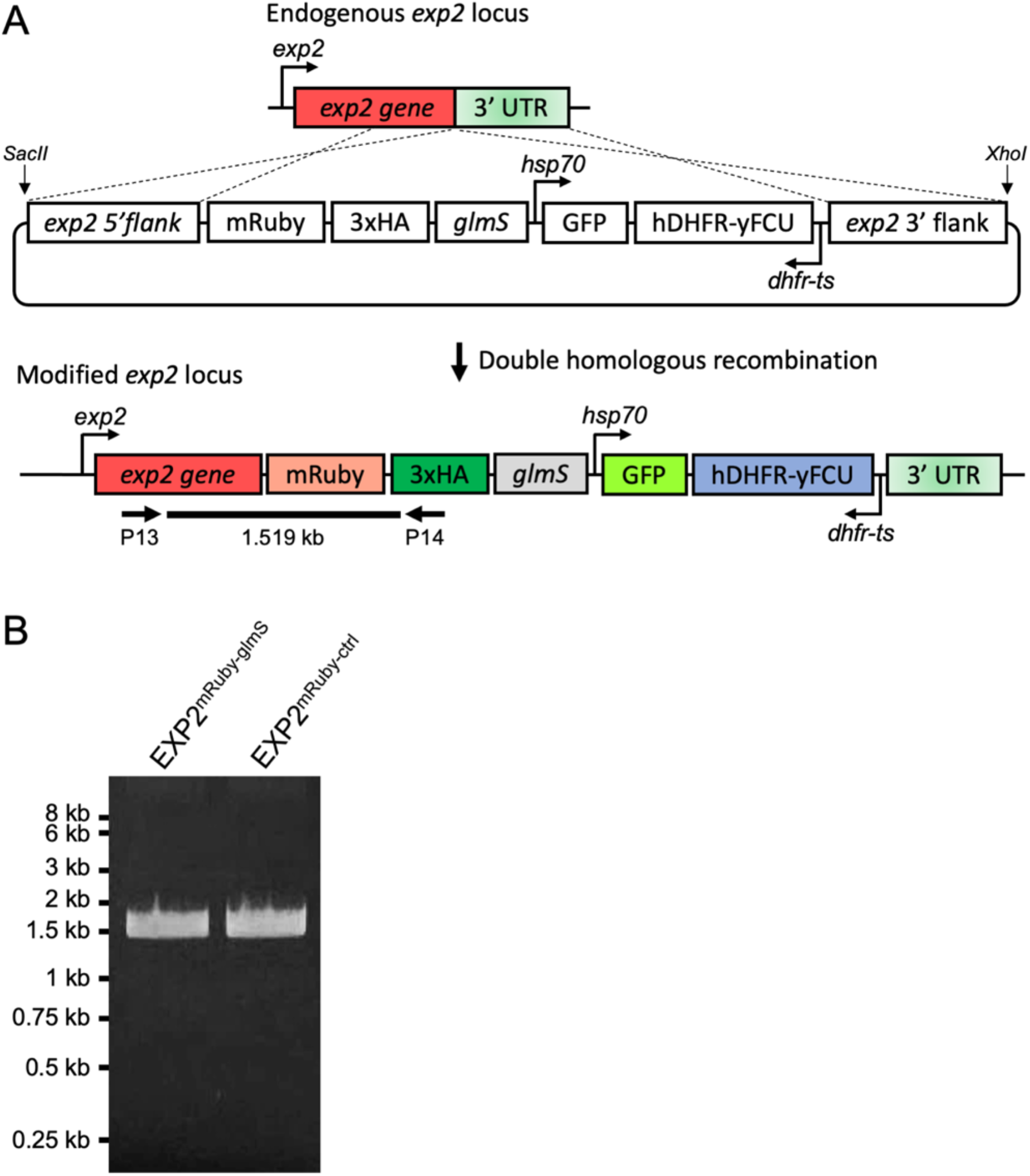
Generation of EXP2-mRuby-3xHA *glmS* parasites. A) Detailed schematic showing modification of the endogenous *exp2* locus by double homologous recombination resulting in a C-terminal mRuby3-3xHA fusion followed by the *glmS* ribozyme immediately downstream of the stop codon in the 3’ UTR. The plasmid also contains a downstream cassette for expression of GFP under the control of the *hsp70* promoter. In parallel, a control parasite line was generated with a plasmid lacking the *glmS* sequence but otherwise identical. B) Diagnostic PCR with primers indicated in the schematic showing successful integration at the *exp2* locus.

**Figure S2.**
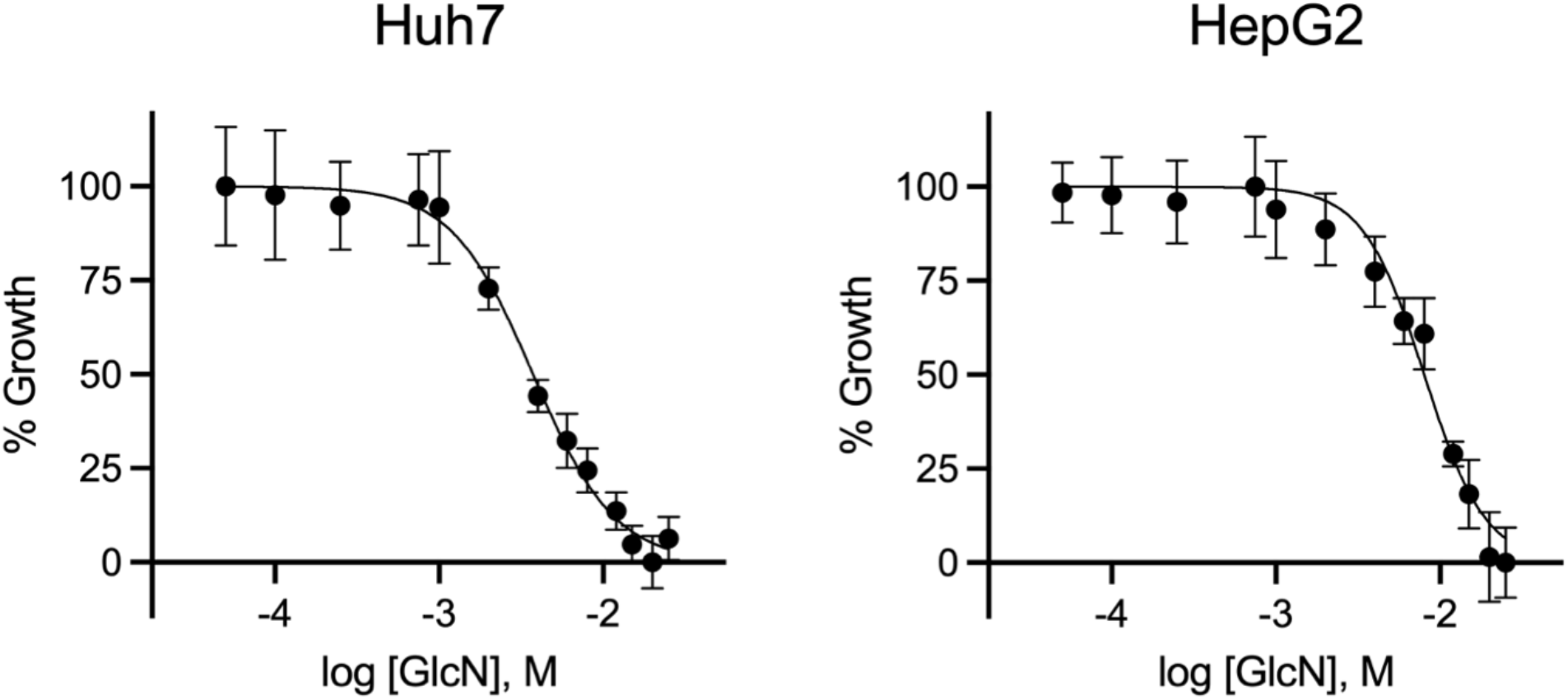
Impact of GlcN on hepatoma cell viability. Dose response curves of Huh7 and HepG2 hepatoma cells measured using a resazurin assay. Curves were fit to the mean of four independent biological replicates, each performed in technical triplicate. Error bars indicate SEM. The EC_50_ was determined to be 3.7 mM for Huh7 cells and 8 mM for HepG2 cells.

**Figure S3.**
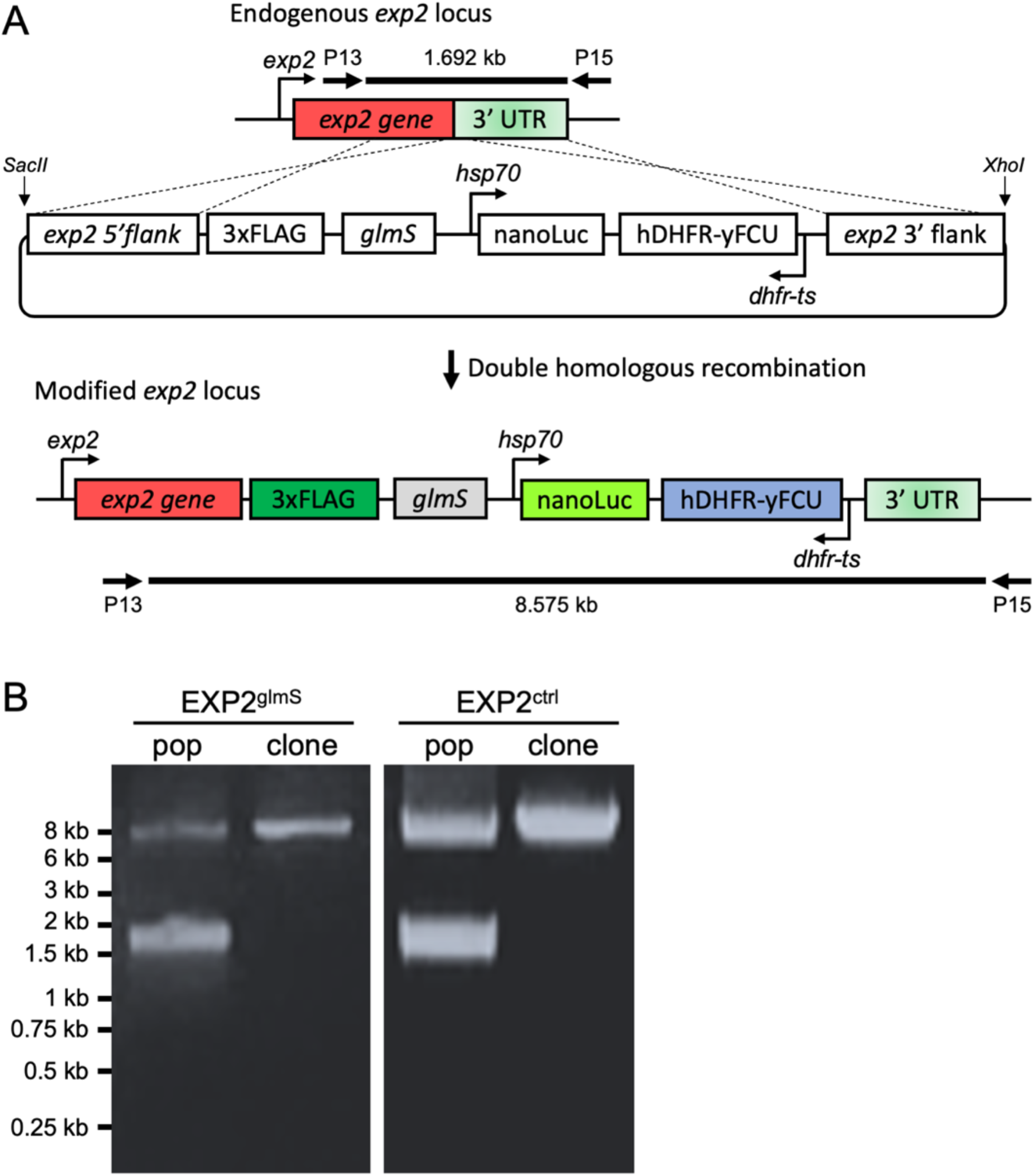
Generation of EXP2^*glmS*^ and EXP2^ctrl^ parasites. A) Schematic showing modification of the endogenous *exp2* locus by double homologous recombination resulting in a C-terminal 3xFLAG fusion followed by the *glmS* ribozyme immediately downstream of the stop codon in the 3’ UTR. The plasmid also contains a downstream cassette for expression of nanoluciferase under the control of *hsp70* promoter. In parallel, a control parasite cell line was generated with a plasmid lacking the *glmS* sequence but otherwise identical. B) Diagnostic PCR with primers indicated in the schematic showing successful integration at the *exp2* locus in transfected populations and absence of the unmodified locus in isogenic clonal populations.

**Figure S4.**
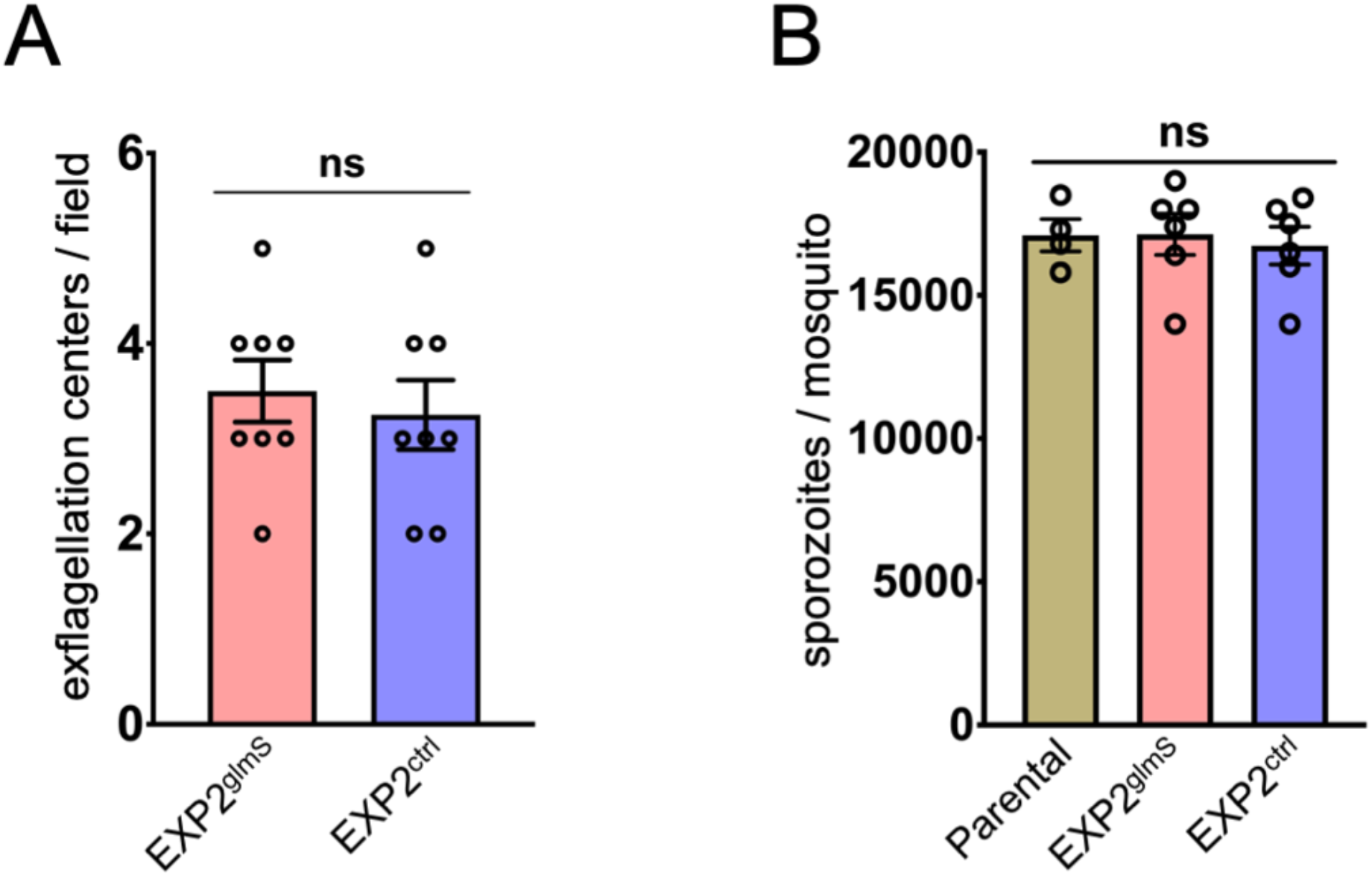
Sporozoite production is unaltered in EXP2^*glmS*^ and EXP2^ctrl^ parasites. A) Quantification of exflagellation centers 5 days post infection with EXP2^*glmS*^ and EXP2^ctrl^ parasites. B) Quantification of sporozoites obtained per mosquito following dissection 21 days post blood feeding for indicated parasite lines. Data points represent independent biological replicates. Error bars indicate SEM.

**Figure S5.**
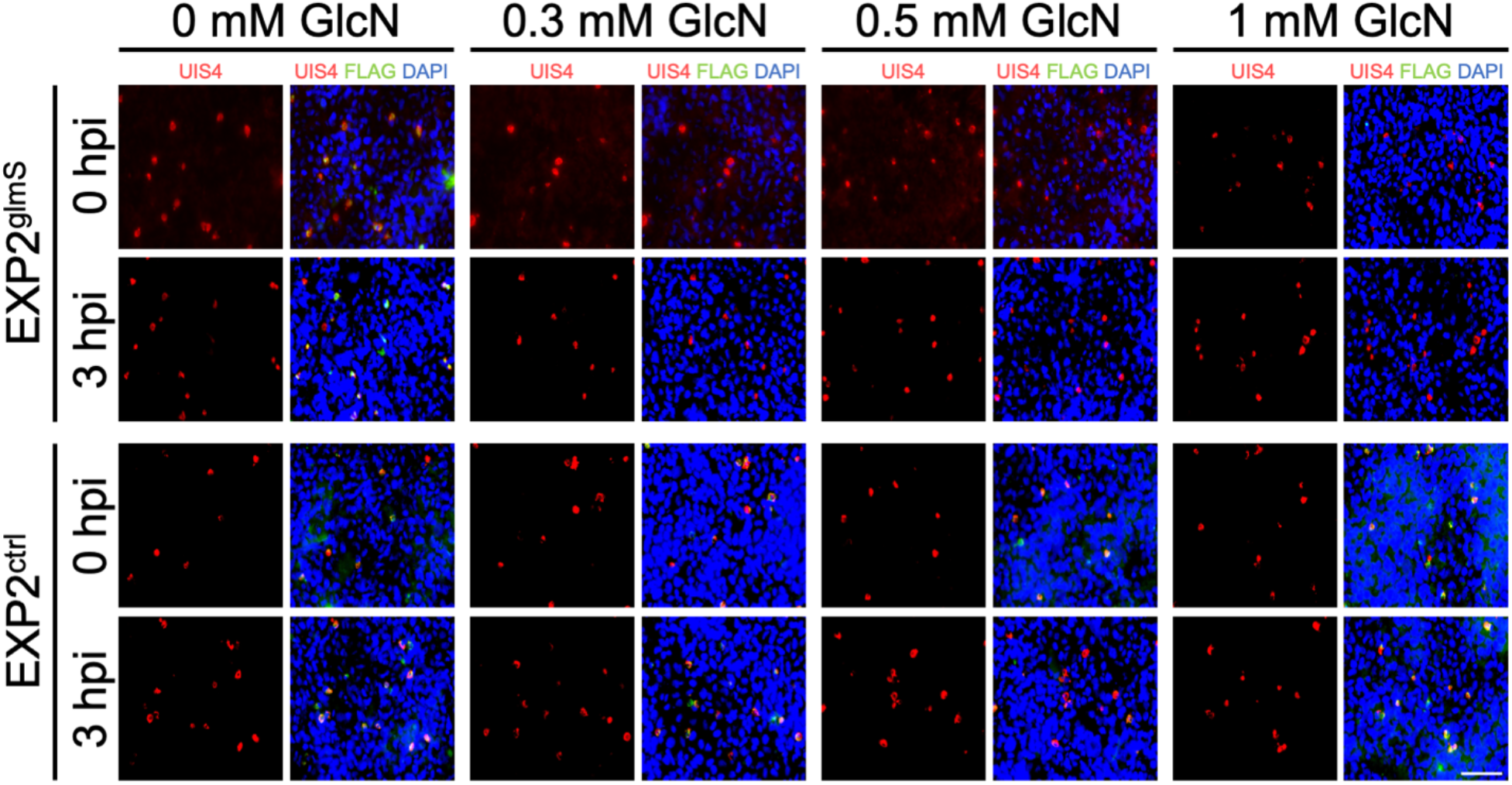
GlcN treatment does not impact EXP2^glmS^ sporozoite invasion. Sporozoites from EXP2^glmS^ parasites were allowed to infect Huh7 cells in the presence or absence of 0.5mM GlcN for 3 hours before monolayers were washed, fixed and processed for IFA. Images were acquired on an automated Keyence microscope and UIS4-positive vacuoles per mm^2^ were quantified. Data from 3 independent biological replicates are shown. Error bars indicate SEM.

**Figure S6.**
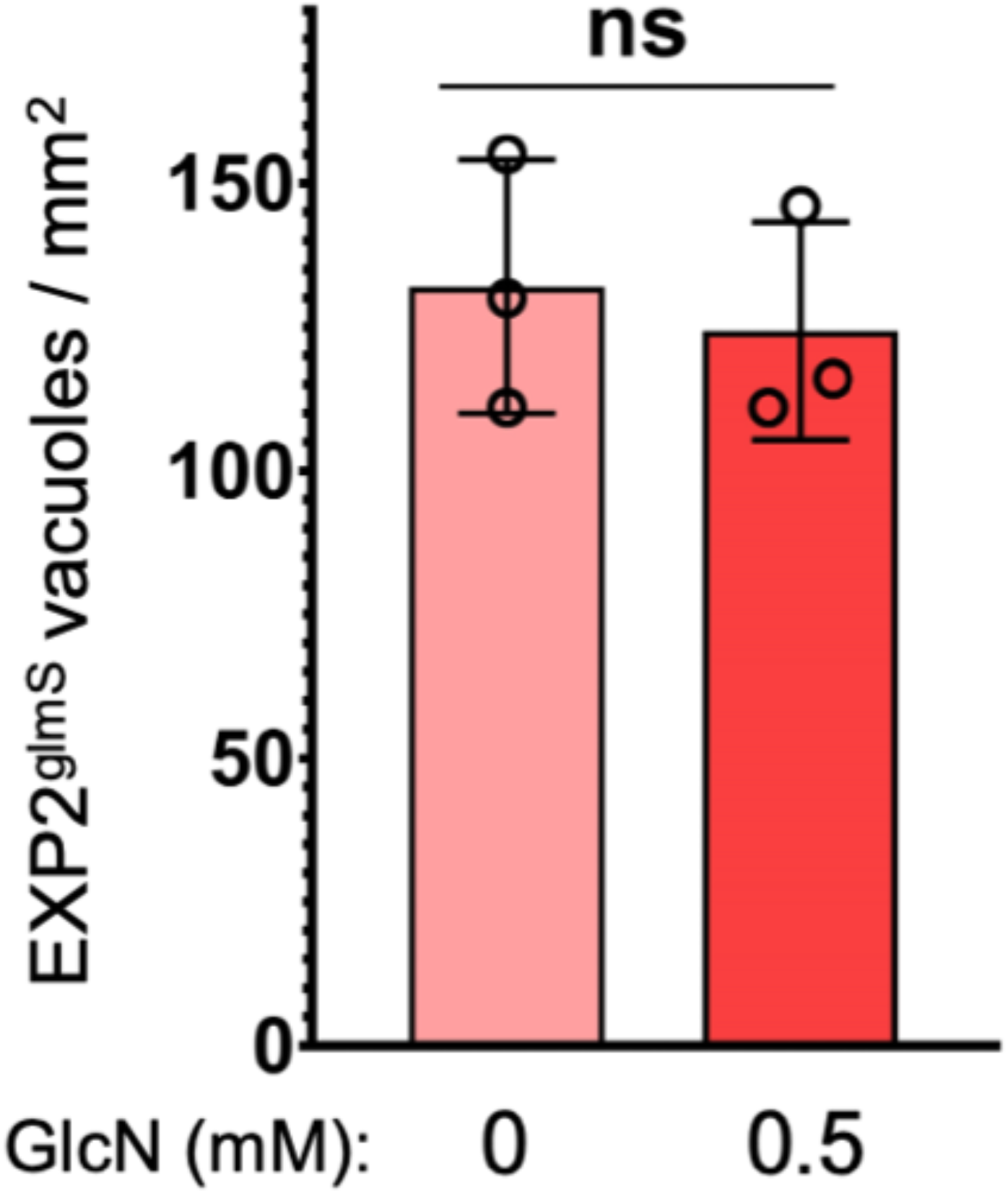
Representative images used for quantification of vacuolar area. Sporozoites were obtained by mosquito dissection and allowed to infect Huh7 monolayers for 3 hours before media change to remove remaining extracellular parasites. GlcN was added at indicated concentrations from the time sporozoites were introduced to Huh7 cultures (0 hpi) or following initial media change (3 hpi). Media was changed daily and cells were fixed and processed for IFA at 48 hpi. Images were acquired on an automated Keyence microscope using a 40X object and analyzed to determine vacuolar area using UIS4 to mark the PVM. Representative fields are shown. Scale bar = 62.5 μm.

**Figure S7.**
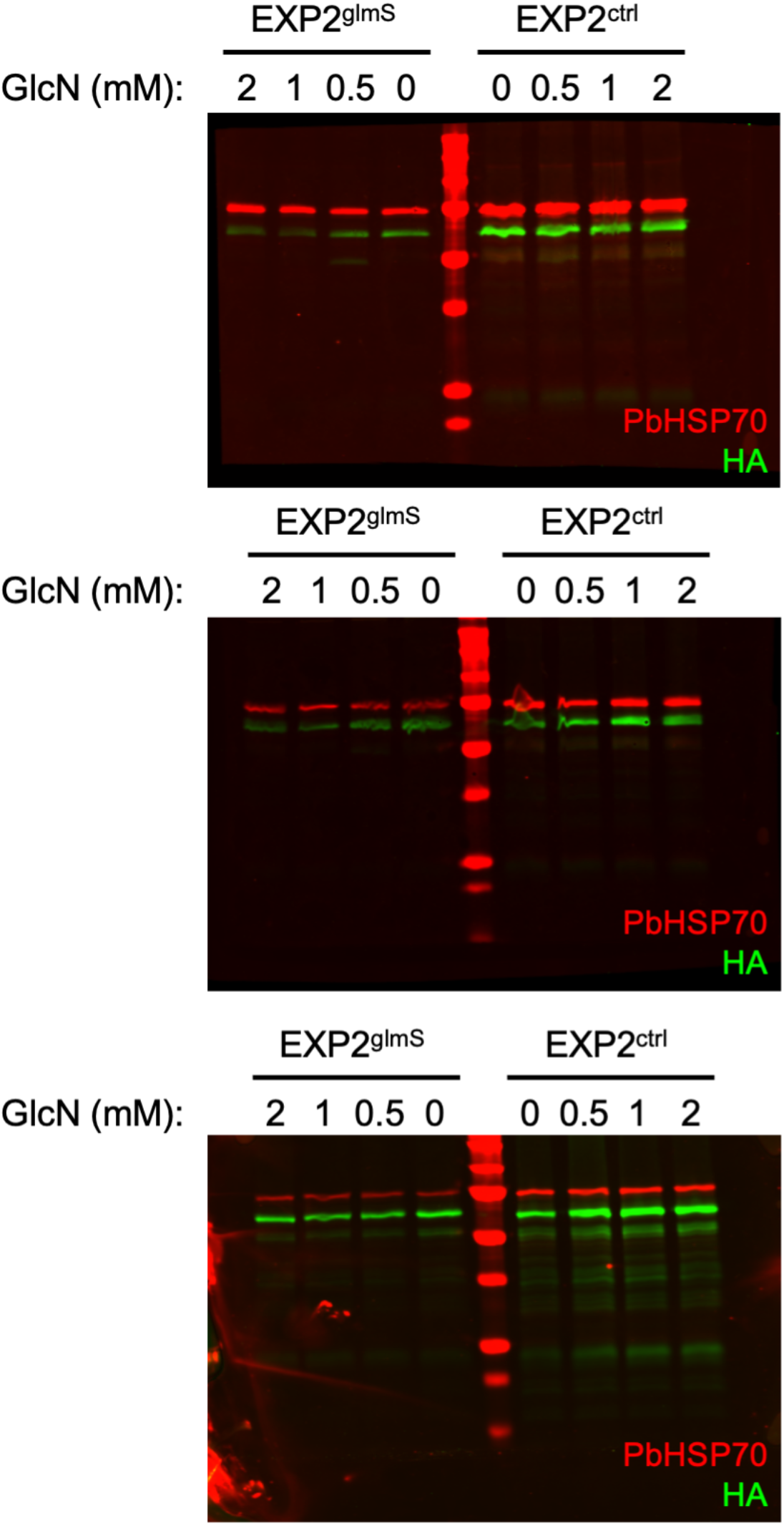
Uncropped Western blots for quantification of EXP2 knockdown in *ex vivo* blood stage experiments in Figure 1.

---

**Table S1.**
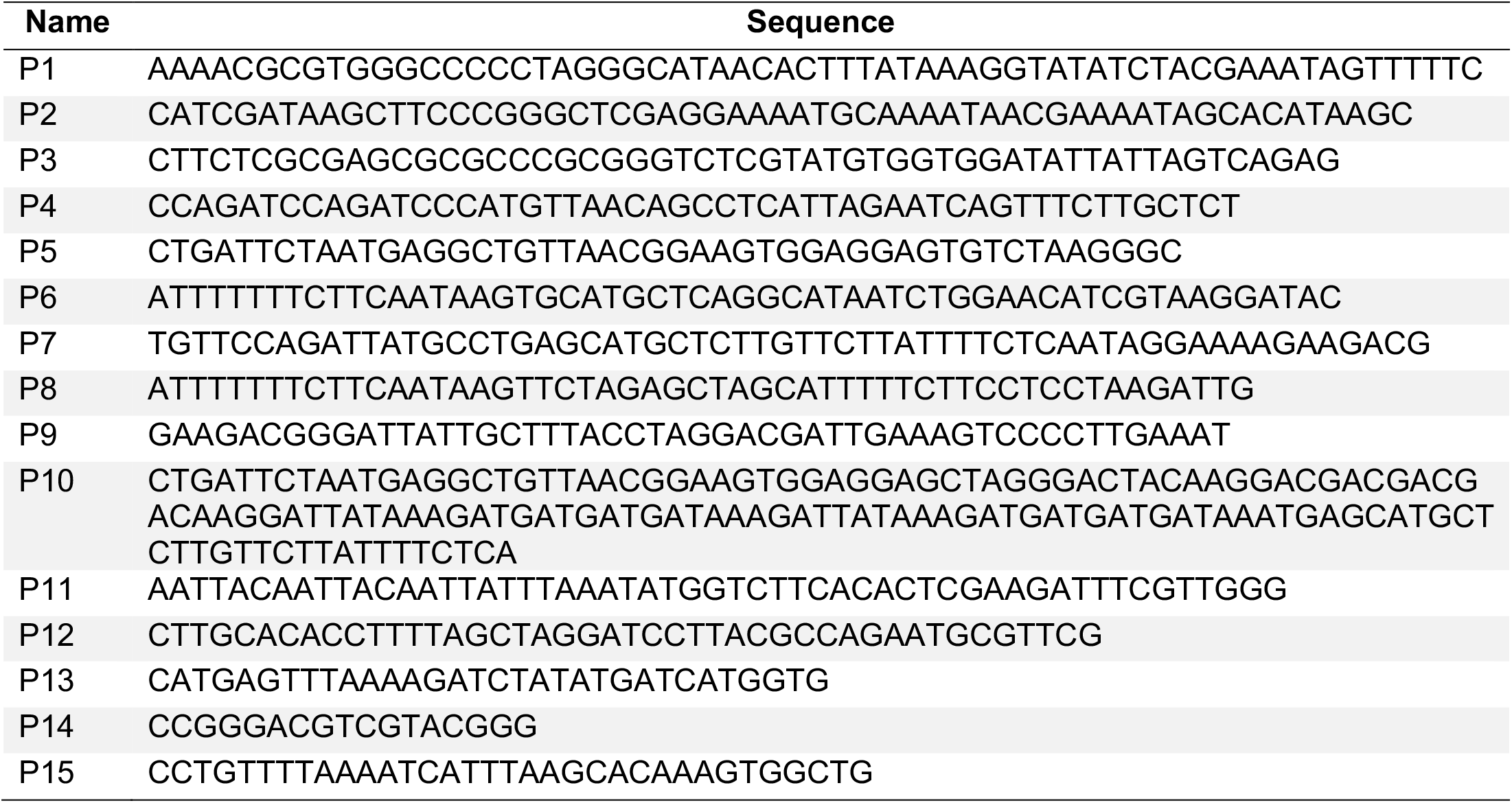
Sequences of primers used in this study.

## Notes

### Competing Interest Statement

The authors have declared no competing interest.

