## Supplemental Figures for "The PTEX pore component EXP2 is important for intrahepatic development during the *Plasmodium* liver stage"

**Figure S1**

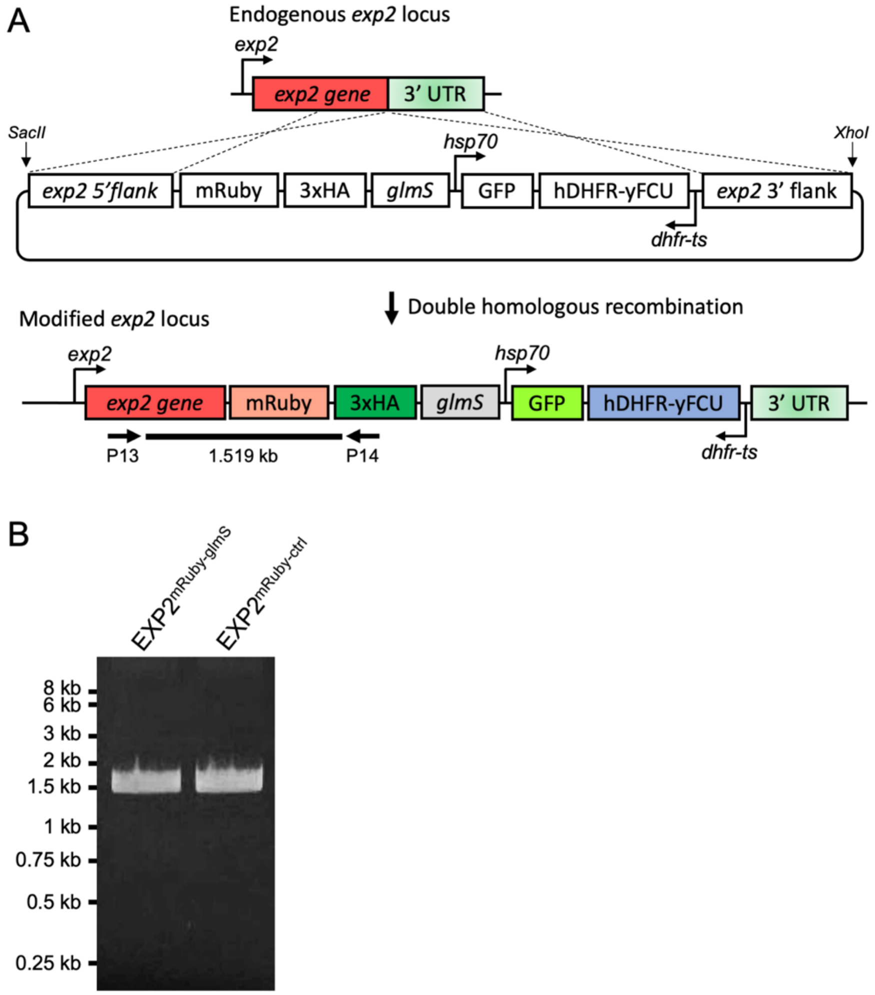
A
Endogenous exp2 locus
exp2
exp2 gene
3' UTR
SacII
XhoI
hsp70
exp2 5'flank
mRuby
3xHA
glmS
GFP
hDHFR-yFCU
exp2 3' flank
dhfr-ts
Double homologous recombination
Modified exp2 locus
exp2
hsp70
exp2 gene
mRuby
3xHA
glmS
GFP
hDHFR-yFCU
3' UTR
P13
1.519 kb
P14
dhfr-ts
B
EXP2mRuby-glmS
EXP2mRuby-ctrl
8 kb
6 kb
3 kb
2 kb
1.5 kb
1 kb
0.75 kb
0.5 kb
0.25 kb

**Figure S2**

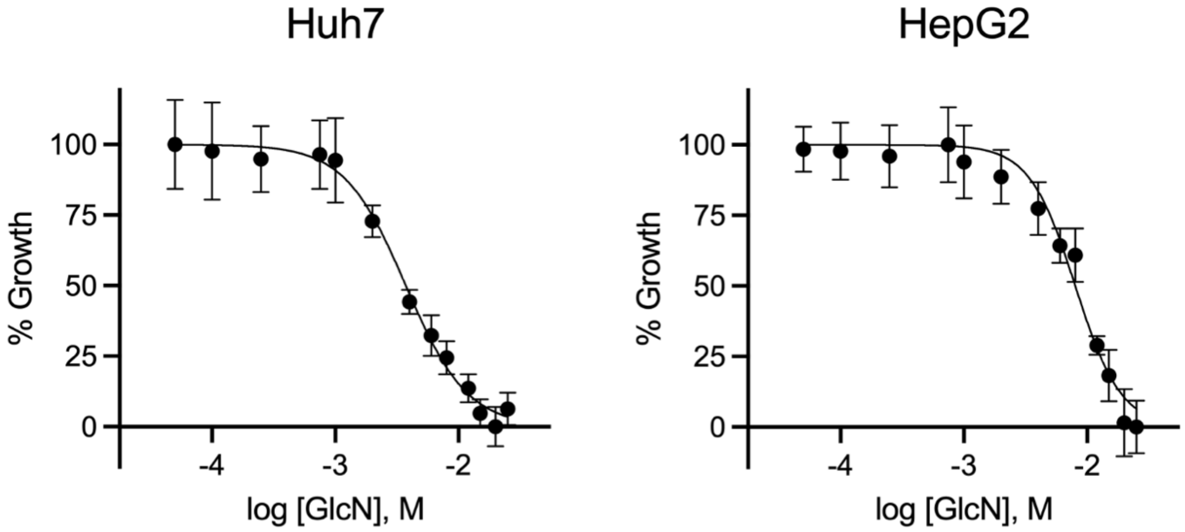

**Figure S3**

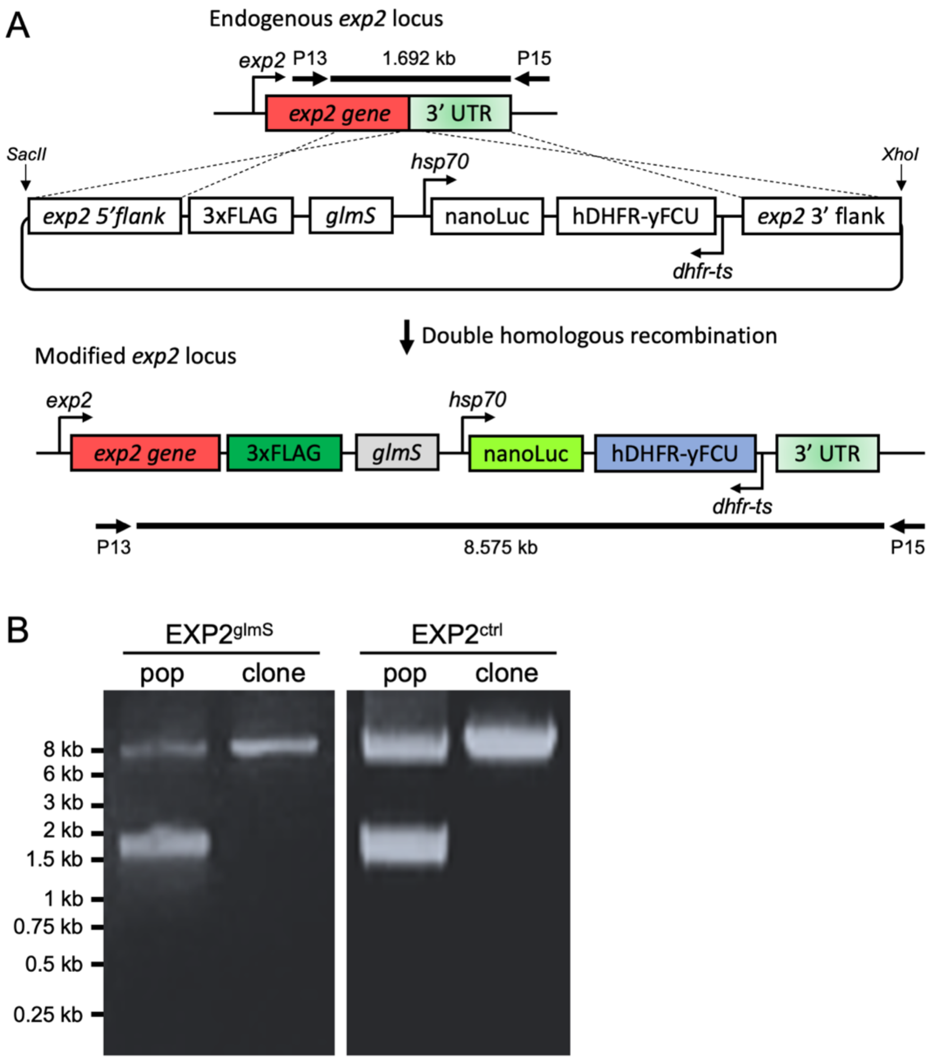
A
Endogenous exp2 locus
exp2
P13
1.692 kb
P15
exp2 gene
3' UTR
SacII
XhoI
hsp70
exp2 5'flank
3xFLAG
glmS
nanoLuc
hDHFR-yFCU
exp2 3' flank
dhfr-ts
Double homologous recombination
Modified exp2 locus
exp2
hsp70
exp2 gene
3xFLAG
glmS
nanoLuc
hDHFR-yFCU
3' UTR
dhfr-ts
P13
8.575 kb
P15
B
EXP2glmS
EXP2ctrl
pop
clone
pop
clone
8 kb
6 kb
3 kb
2 kb
1.5 kb
1 kb
0.75 kb
0.5 kb
0.25 kb

**Figure S4**

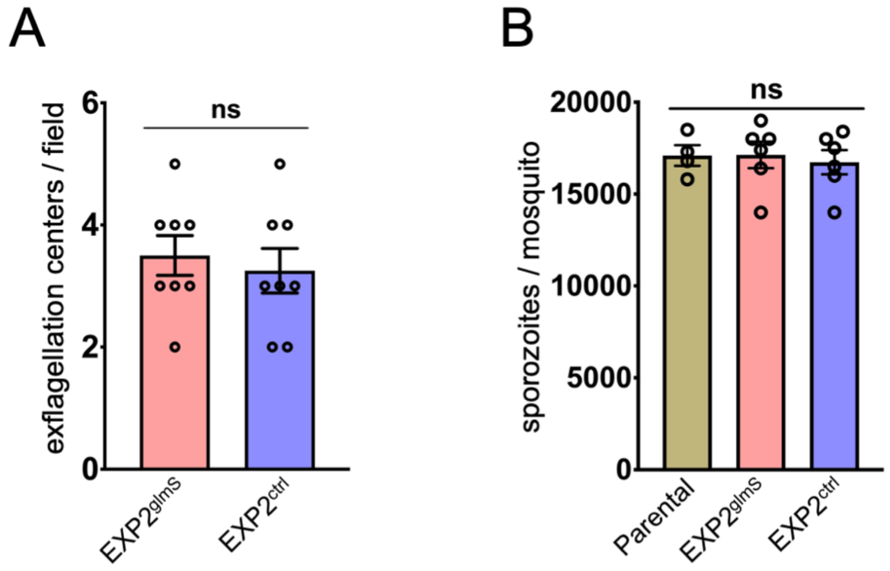
A
B
ns
ns
exflagellation centers / field
6
4
2
0
EXP2glmS
EXP2ctrl
sporozoites / mosquito
20000
15000
10000
5000
0
Parental
EXP2glmS
EXP2ctrl

**Figure S5**

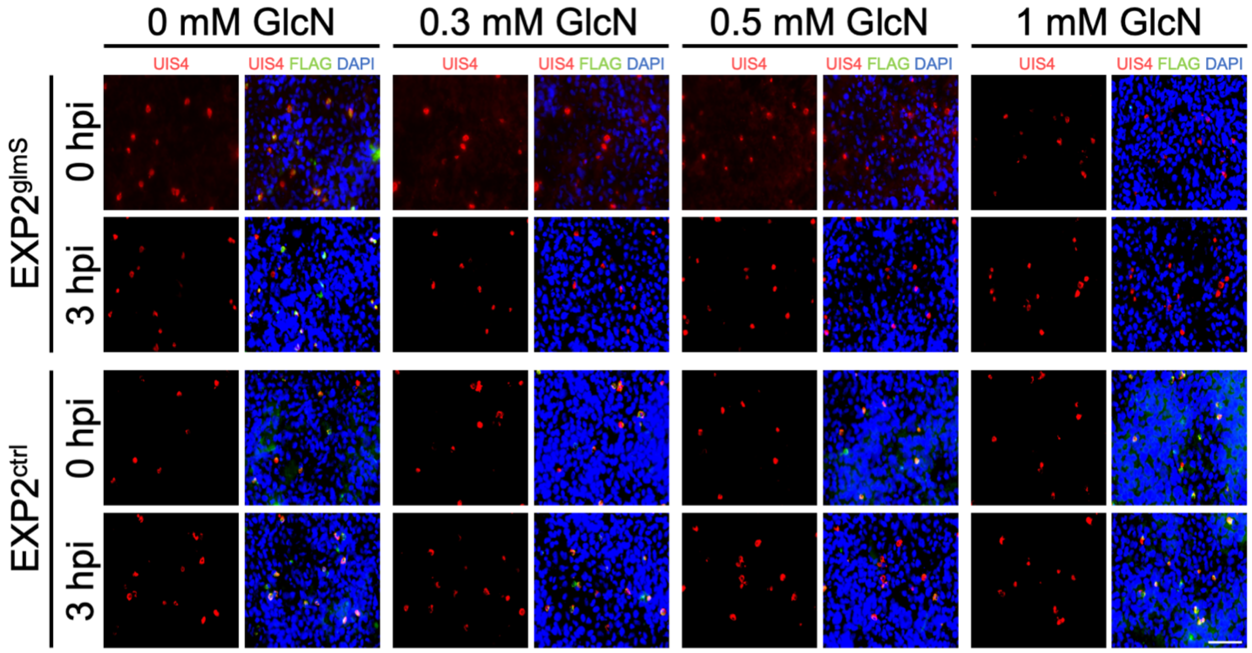
0 mM GlcN
0.3 mM GlcN
0.5 mM GlcN
1 mM GlcN
UIS4
UIS4 FLAG DAPI
EXP2glmS
EXP2ctrl
0 hpi
3 hpi

**Figure S6**

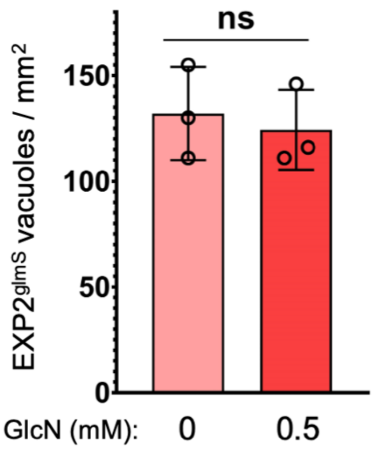

Figure S7

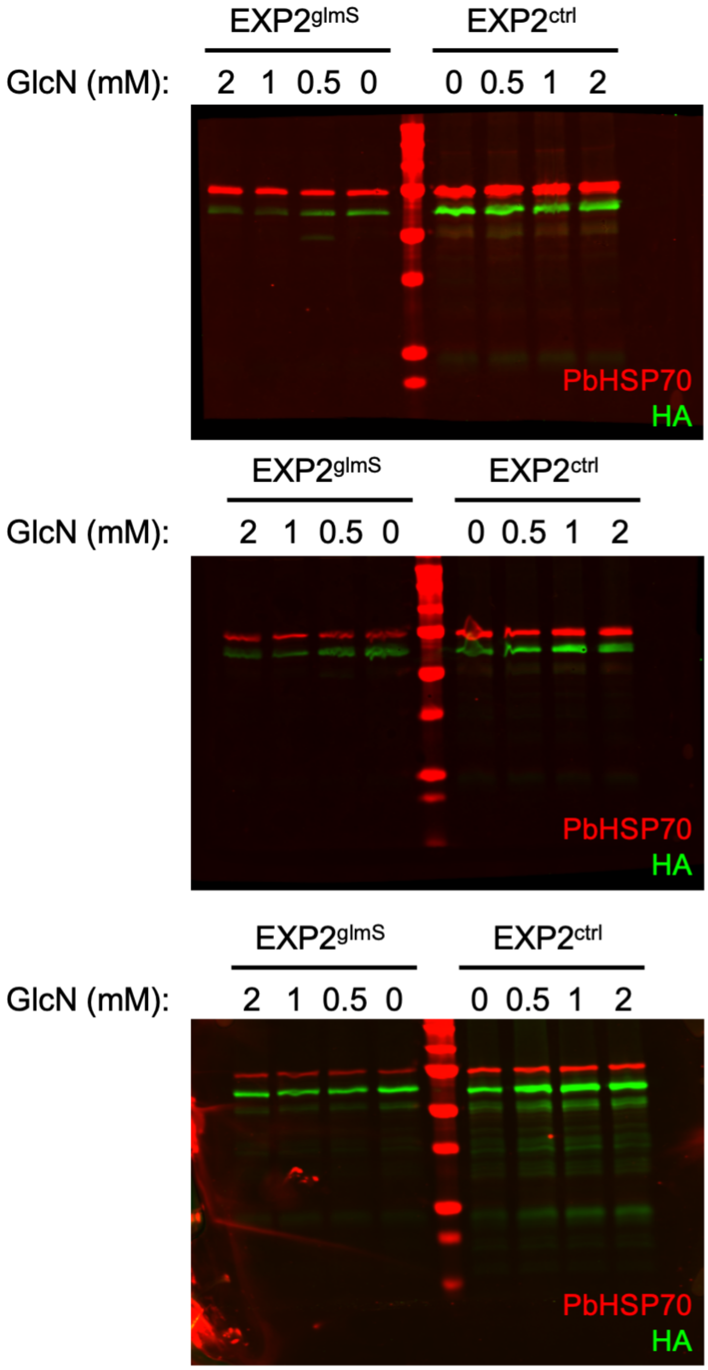
EXP2glmS
EXP2ctrl
GlcN (mM):
2 1 0.5 0
0 0.5 1 2
PbHSP70
HA
EXP2glmS
EXP2ctrl
GlcN (mM):
2 1 0.5 0
0 0.5 1 2
PbHSP70
HA
EXP2glmS
EXP2ctrl
GlcN (mM):
2 1 0.5 0
0 0.5 1 2
PbHSP70
HA

**Table S1.** Sequences of primers used in this study.

| Name | Sequence |
|---|---|
| P1 | AAAACGCGTGGGCCCCCTAGGGCATAACACTTTATAAAGGTATATCTACGAAATAGTTTTTC |
| P2 | CATCGATAAGCTTCCCGGGCTCGAGGAAAATGCAAAATAACGAAAATAGCACATAAGC |
| P3 | CTTCTCGCGAGCGCGCCCGCGGGTCTCGTATGTGGTGGATATTATTAGTCAGAG |
| P4 | CCAGATCCAGATCCCATGTTAACAGCCTCATTAGAATCAGTTTCTTGCTCT |
| P5 | CTGATTCTAATGAGGCTGTTAACGGAAGTGGAGGAGTGTCTAAGGGC |
| P6 | ATTTTTTTCTTCAATAAGTGCATGCTCAGGCATAATCTGGAACATCGTAAGGATAC |
| P7 | TGTTCCAGATTATGCCTGAGCATGCTCTTGTTCTTATTTTCTCAATAGGAAAAGAAGACG |
| P8 | ATTTTTTTCTTCAATAAGTTCTAGAGCTAGCATTTTTCTTCCTCCTAAGATTG |
| P9 | GAAGACGGGATTATTGCTTTACCTAGGACGATTGAAAGTCCCCTTGAAAT |
| P10 | CTGATTCTAATGAGGCTGTTAACGGAAGTGGAGGAGCTAGGGACTACAAGGACGACGACG ACAAGGATTATAAAGATGATGATGATAAAGATTATAAAGATGATGATGATAAATGAGCATGCT CTTGTTCTTATTTTCTCA |
| P11 | AATTACAATTACAATTATTTAAATATGGTCTTCACACTCGAAGATTTCGTTGGG |
| P12 | CTTGCACACCTTTTAGCTAGGATCCTTACGCCAGAATGCGTTCG |
| P13 | CATGAGTTTAAAAGATCTATATGATCATGGTG |
| P14 | CCGGGACGTCGTACGGG |
| P15 | CCTGTTTTAAAATCATTTAAGCACAAAGTGGCTG |
